# Modelling motion-induced signal corruption in steady-state diffusion MRI

**DOI:** 10.1101/2025.08.19.670858

**Authors:** Benjamin C. Tendler, Wenchuan Wu, Karla L. Miller, Aaron T. Hess

## Abstract

**Purpose:** Diffusion-weighted steady-state free precession (DW-SSFP) is a diffusion imaging sequence achieving high SNR efficiency. A key challenge for in vivo DW-SSFP is the sequence’s severe motion sensitivity, currently limiting investigations to low or no motion regimes. Here we establish a framework to both (1) model and (2) correct for the impact of subject motion associated with the underlying magnetisation distribution of DW-SSFP.

**Theory and Methods:** An extended phase graphs (EPG) representation of the 1D DW-SSFP signal was established incorporating a motion operator describing rigid body and pulsatile motion. The representation was validated using Monte Carlo simulations, and subsequently integrated into a data fitting routine for motion estimation and correction. The fitting routine was evaluated using both simulations and a voxelwise correction applied to in vivo experimental 2D low-resolution single-shot timeseries DW-SSFP data acquired in the human brain in three healthy volunteers, with a tensor reconstructed from the motion-corrected experimental DW-SSFP data.

**Results:** The proposed EPG-motion framework gives excellent agreement to complementary Monte Carlo simulations, demonstrating that diffusion coefficient estimation is robust over a range of motion and SNR regimes. Tensor estimates from the motion-corrected experimental DW-SSFP data give good visual agreement to complementary diffusion-weighted spin-echo (DW-SE) data acquired in the same subject, considerably reducing orientation-dependent motion-induced biases.

**Conclusion:** Temporal information capturing the evolution of the DW-SSFP signal can be used to retrospectively (1) estimate subject motion and (2) reconstruct motion-corrected DW-SSFP data. Open-source software is provided, facilitating future investigations into the impact of subject-motion on DW-SSFP acquisitions.

## Introduction

In diffusion MRI, applied diffusion encoding gradients sensitise signals to the microscopic (μm) scale, with diffusion processes in tissue leading to attenuation of signal magnitude^1^. Measured signals are also intrinsically sensitive to subject motion, with small tissue displacements (e.g. arising from bulk, pulsatile, or respiratory induced motion) leading to changes in signal phase^2,3^. The presence of motion-induced artefacts in reconstructed images is strongly dependent on the incorporated readout scheme. The combination of phase-inconsistent k-space data associated with segmented readouts can give rise to considerable artefacts in reconstructed magnitude images^4^, impacting the quantification of diffusion attenuation. Diffusion-weighted spin-echo (DW-SE)^1^ sequences incorporating single-shot readouts are typically considered motion robust^5,6^, with a notable exception of motion induced ‘dropout’ arising when the echo is displaced from the k-space field of view (FOV)^7–9^.

Diffusion-weighted steady-state free precession (DW-SSFP)^10–13^ is an alternative to the DW-SE for diffusion MRI investigations. DW-SSFP is typically implemented as a 3D sequence, with a diffusion encoding module consisting of a single diffusion gradient and RF pulse per TR (Figure 1). The signal-forming mechanisms of DW-SSFP have previously demonstrated high-SNR efficiency^14,15^, with the sequences’ short TR (∼20 to 40 ms) compatible with a segmented readout acquiring a single line (or a few lines) of k-space per TR, leading to images with minimal geometric distortions. Whilst offering considerable potential for in vivo investigations, previous work has demonstrated that the DW-SSFP acquisitions are highly sensitive to both rigid body (head translation/rotation) and pulsatile (i.e. arising from the cardiac cycle) motion^16–21^. At present, the motion-sensitivity of DW-SSFP limits investigations to low (cartilage^15,22^, peripheral nerves^23^) or no (post-mortem^14^) motion imaging regimes.

**Figure 1:**
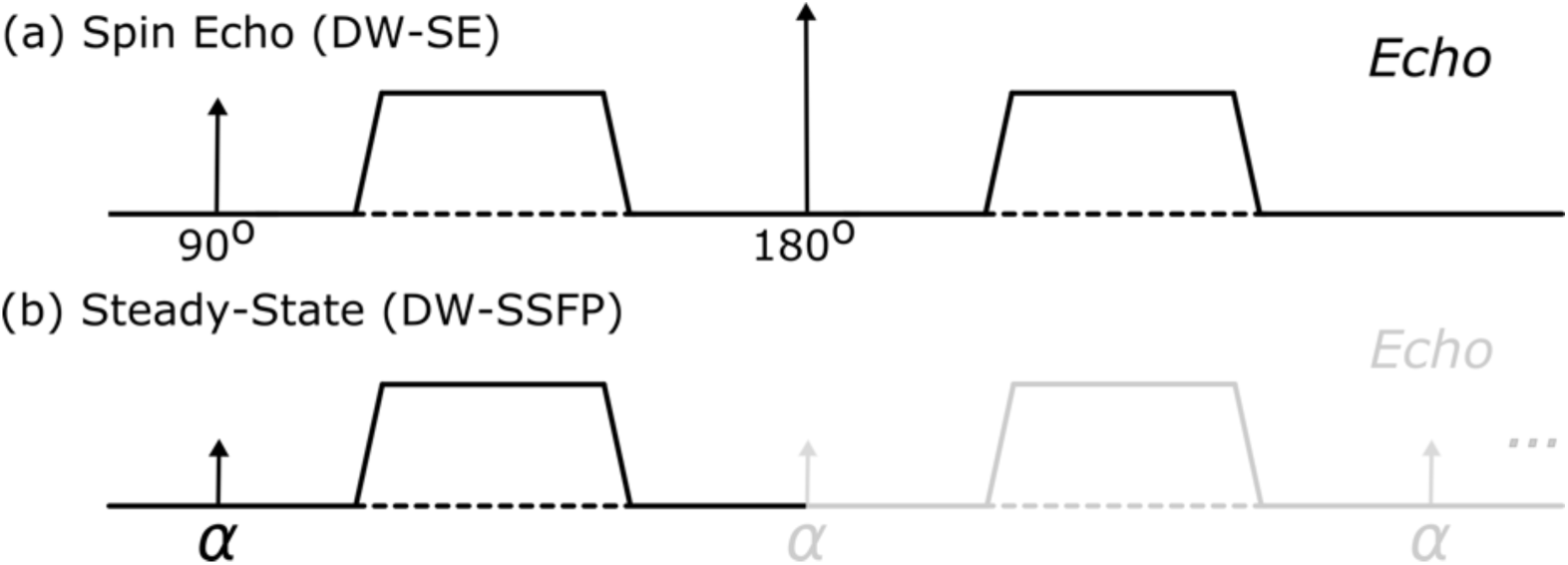
Diffusion encoding. Diffusion encoding modules of the (a) diffusion-weighted spin-echo (DW-SE) and (b) diffusion-weighted steady-state free precession (DW-SSFP) sequence. DW-SSFP consists of a single RF pulse and diffusion gradient per TR (typically 20 – 40 ms). Magnetisation accumulates diffusion contrast over several TRs, consistent with dephasing and rephasing of magnetisation due to gradient pairs in the DW-SE. The steady-state DW-SSFP signal (reached after many TRs) can be considered a superposition of multiple magnetisation components with different histories – e.g., pairs of diffusion-weighted gradients separated by a variable integer number of TRs that lead to different degrees of diffusion weighting. The measured signal is the sum of these different magnetisation components, with diffusion-weighting dependent on the applied diffusion gradient, TR, flip angle, T1, T2, and diffusion coefficient^32,33^.

Several motion-correction methods have been previously established to address motion artefacts arising from segmented diffusion MRI acquisitions. Typically these methods introduce navigator echoes^24,25^ or ‘self-navigating’ readouts (e.g. PROPELLER^26,27^, SNAILS^28,29^) to the sequence, or exploit sensitivity encodings (e.g. MUSE^30^, MUSSELS^31^) to estimate and correct for motion-induced phase inconsistencies across acquired k-space segments. These techniques have demonstrated considerable promise for the DW-SE sequence, raising a simple question – can we directly translate an existing method to correct for subject motion in DW-SSFP?

Unfortunately, a key challenge for DW-SSFP is the specialised nature of how motion corrupts the signal associated with the underlying magnetisation distribution (i.e. independent of the readout scheme and k-space coverage). Specifically, whilst subject motion corrupts the DW-SE signal phase, in DW-SSFP subject motion corrupts both the signal phase *and* magnitude^3^ and persists over many TRs (Figure 2). Whilst conventional motion-correction techniques can be incorporated to attenuate artefacts associated with the sequence’s segmented readout^16–20^, we require further extensions to account for the signal forming mechanisms of DW-SSFP.

**Figure 2:**
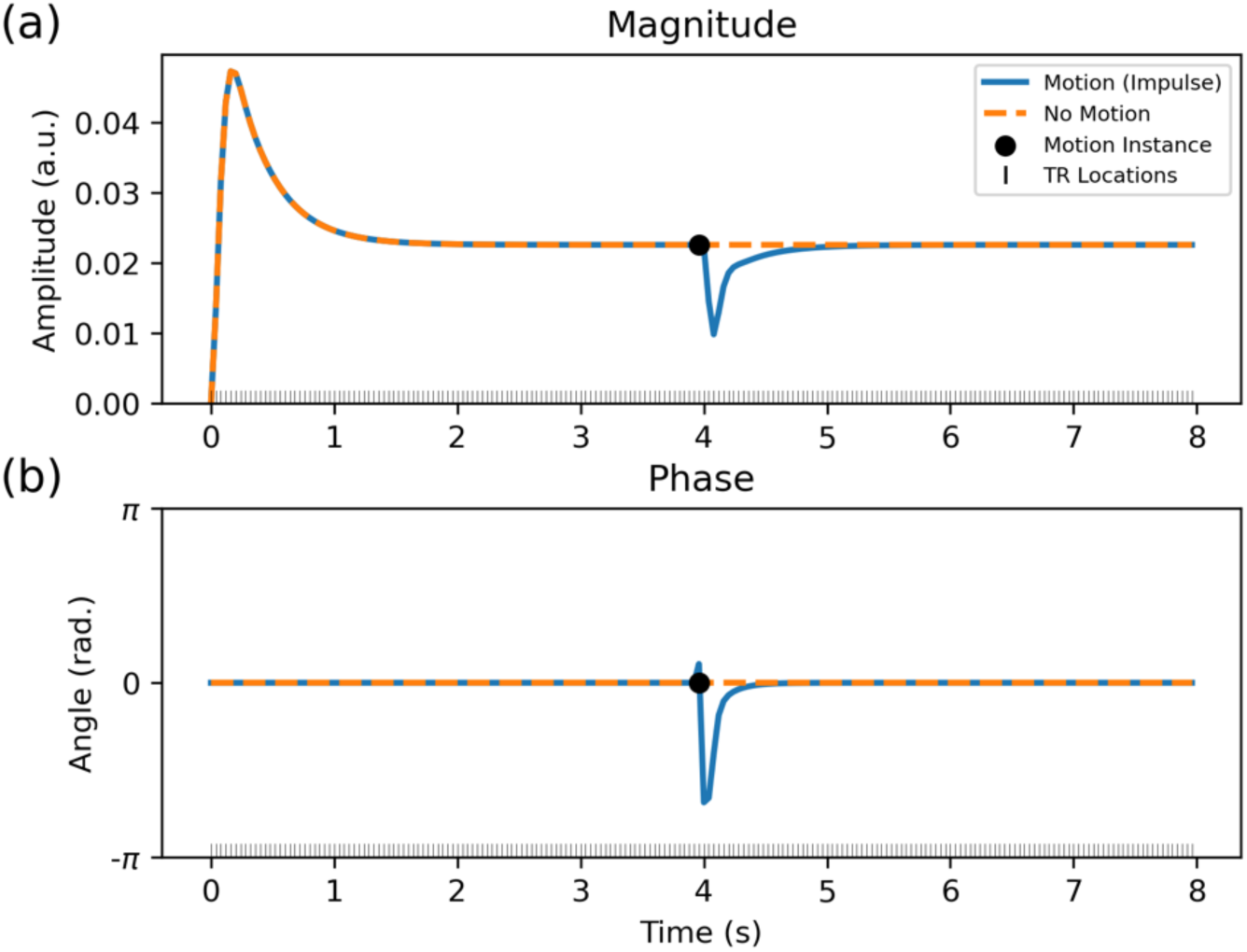
DW-SSFP signal evolution with impulse motion. Simulated 1D time-series profile of the DW-SSFP (a) magnitude and (b) phase signal over 200 TRs. After ∼50 TRs (2 s), the signal reaches a steady state. Impulse motion simulated during a single TR (black dot) perturbs the steady state. This motion-corrupted magnetisation persists and contributes to future TRs, leading to a slow recovery to the original steady state, with both the magnitude and phase signal impacted. Sequence parameters based on the experimental investigation performed in Miller & Pauly^17^, setting G = 40 mT/m, δ = 6.5 ms, α = 30°, TR = 40 ms and 𝛟 = 0°. Impulse motion was modelled using a velocity of 1.5 mm/s along the diffusion gradient orientation during a single TR (40 ms) using the proposed EPG-motion framework (see Theory), leading to a total displacement of 0.06 mm. Sample parameters based on approximate in vivo values in brain tissue at 3T^35^, defining D = 1 ⋅ 10^−3^ mm^2^/s, T_1_ = 832 ms and T_2_ = 110 ms.

Specifically, as DW-SSFP is a steady-state sequence, magnetisation excited in a single TR persists and evolves over multiple TRs. Over time the DW-SSFP signal reaches a steady-state (Figure 2), with the signal reflecting a weighted sum of magnetisation components with different evolution histories.

Motion at any timepoint leads to phase corruption of the excited magnetisation, which over future TRs will combine with magnetisation components with different motion histories and associated phase corruption, leading to phase cancellation and subsequent magnitude loss (Figure 2). This includes the combination of phase corrupted magnetisation with ‘fresh’ magnetisation (i.e. longitudinally recovered magnetisation that has been excited into the transverse plane) that has no motion history. This form of magnitude loss is independent of the readout scheme and k-space coverage. From the perspective of Extended Phase Graphs^34^ (EPG), magnitude corruption of the DW-SSFP signal arises from the combination of magnetisation pathways with different phase values during the evolution of the phase graph.

Several methods have been proposed to address motion artefacts in DW-SSFP^16–20^, generally by adopting navigator-based phase corrections that were originally proposed for DW-SE. A key limitation of these previous methods is that they do not account for the loss of signal magnitude arising from the combination of magnetisation components with different evolution histories, leading to residual artefacts in reconstructed images. One notable exception is work by O’Halloran et al.^21^ establishing a prospective motion-correction method for DW-SSFP utilising real-time information to estimate and correct for deviations in image phase arising from rigid-body motion of the subject, providing partial attenuation of magnitude signal loss.

In this work, we perform a theoretical investigation, and experimental validation, into the impact of subject motion on the DW-SSFP signal associated with the underlying magnetisation distribution (i.e. independent of the readout and k-space coverage). We demonstrate that the loss of motion-induced signal magnitude associated with the magnetisation distribution can be estimated and corrected *retrospectively* using timeseries data capturing the evolution of the DW-SSFP signal. To achieve this, we first introduce a theoretical framework based on EPG^34^ (building on a previous abstract^36^) to characterise how the 1D DW-SSFP signal is corrupted by motion arising from (1) rigid-body motion and (2) brain pulsatility associated with the cardiac cycle. We validate the proposed framework using Monte Carlo simulations, and use the framework to visualise how changes in the magnetisation distribution arising from subject motion impacts DW-SSFP data. We demonstrate that motion-corrupted DW-SSFP data can contain sufficient information to estimate and reconstruct motion-corrected estimates, evaluating the framework experimentally by applying a voxelwise correction to in vivo 2D low-resolution single-shot DW-SSFP timeseries data acquired in the human brain.

Our work addresses a specific sub-problem of motion corruption arising from the magnetisation distribution of DW-SSFP under the assumption an instantaneous readout (i.e. all k-space data required to form an image is acquired in a single TR, here approximated experimentally using a single-shot readout). The proposed model forms the foundation for future DW-SSFP motion-correction methods incorporating information about the k-space readout and coverage, required for translation to a conventional 3D multi-shot DW-SSFP acquisition. Open-source software is provided to enable researchers to replicate many of the findings in this manuscript and perform their own investigations on the impact of subject motion on DW-SSFP data (github.com/BenjaminTendler/MotionCorrectionDWSSFP).

## Theory

### Extended Phase Graphs (EPG)

In this work, we use EPG to characterise the impact of motion on the DW-SSFP signal. Briefly, EPG describes a widely adopted framework for modelling the magnetisation evolution of an MRI sequence. It provides a solution to the Bloch equations^37^, reparameterising magnetisation as a Fourier basis with distinct phase states of dephasing order *k*, defining:

- Longitudinal (*Z*^#^) and transverse (*F*^#^) magnetisation components with different *k* states (*Z*^#^_*k*_, *F*^#^_*k*_).
- RF pulses acting as an operator that mixes *Z*^#^_*k*_ and *F*^#^_*k*_ components of a given *k*.
- Magnetic field gradients acting on *F*^#^_*k*_ components by increasing or decreasing their dephasing order.

EPG describes the magnetisation evolution in each TR as a series of steps. An RF pulse operator first mixes *Z*^#^_*k*_ and *F*^#^_*k*_ components, followed by the application of additional operators (describing relaxation, motion, diffusion etc) during the TR, with the dephasing order ( *k* ) of magnetisation components updated based on the applied magnetic field gradients. Mathematically, the EPG framework formalises magnetisation components as 3×1 vectors of *F*^#^_*k*_ and *Z*^#^_*k*_ states, acted on by 3×3 operator matrices. For readers unfamiliar with EPG signal representations, we recommend the review article by Weigel^34^.

### Extended Phase Graphs and diffusion

The impact of diffusion attenuation has been previously incorporated into an EPG framework via a diffusion operator^38^, **D**. For a periodic sequence, in which each TR contains a diffusion gradient of duration 𝛿, **D** is given by:

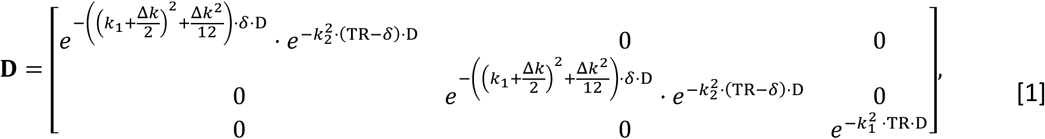

where *k*_1_and *k*_2_ are the dephasing order before and after application of the diffusion gradient, Δ*k* = *k*_2_ − *k*_1_, and *D* is the diffusion coefficient. When considering diffusion gradients, we can define the dephasing order as an integer multiple of the diffusion gradient’s q-value, *q* = ∫ 𝛾 ⋅ *G*(𝑡) ⋅ *dt* , with the change in dephasing order during the application of a diffusion gradient equivalent to Δ*k* = *q*.

The product of the two exponentials present in matrix elements **D**_1,1_ and **D**_2,2_ characterises the diffusion attenuation associated with transverse magnetisation during (1) the application of a diffusion gradient of duration 𝛿 and (2) the remaining TR period. The exponential at location **D**_3,3_characterises the diffusion attenuation experienced by longitudinal magnetisation in a single TR.

### Extended Phase Graphs and motion

Analogous to the description of the diffusion operator in the previous section, we can characterise motion in EPG via a motion operator, 𝐉. In this work, we extend previous descriptions of 𝐉^34^ to incorporate (1) gradients of fixed duration, (2) subject rotation, and (3) time-dependence. Assuming motion as linear (i.e. constant translational/rotational velocity), we can define:

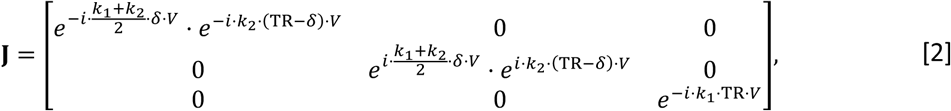

where:

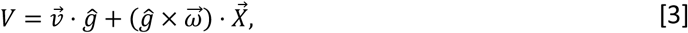

*ĝ*= unit vector describing gradient orientation, 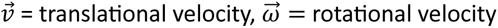 and 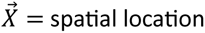 (relative to the centre of rotation). Eq. [2] demonstrates that motion leads to a change in the phase of a magnetisation component, with the polarity of motion (i.e. positive or negative) preserved in the sign of the phase change. To incorporate changes in the motion profile across TRs, we can define:

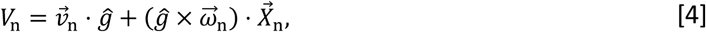

where the subscript indicates the parameters for the n^th^ TR, and 𝑉_n_is a scalar constant representing the component of instantaneous velocity along the orientation of the diffusion gradient during a given TR. We subsequently refer to the evolution of 𝑉_n_ across multiple TRs as 𝑉(𝑡).

The steady-state diffusion signal persists for several seconds over multiple TRs, with the signal lifetime dictated primarily by T^1^. When considering forms of motion that are dependent on the spatial location 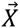(e.g. rotations), the motion operator must account for cumulative changes in 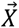 over the time-course of an acquisition. Assuming a single time-step per TR, we define:

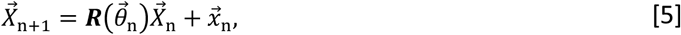

where 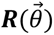 is a 3D rotation matrix operation = 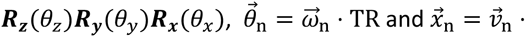 TR. We note that the proposed motion operator is not specific to DW-SSFP, and could be incorporated into EPG schemes investigating the impact of motion on arbitrary MRI sequences, including diffusion-weighted double-echo steady-state (DW-DESS)^39–41^ and MR fingerprinting^42,43^ methods.

As implied from Eq. [3], only velocity components parallel to the applied diffusion gradient *ĝ* contribute to the motion operator in Eq. [2] and lead to a motion-induced signal change. As the velocity profile of rigid body and pulsatile motion in the brain are orientation dependent, DW-SSFP data acquired with different diffusion gradient orientations will be associated with different motion profiles, leading to signal corruption beyond a global loss of SNR. Beyond orientation effects, the degree of signal corruption is dependent on the properties of the diffusion-encoding scheme and the evolution of the motion profile over time (Eq. [4]).

Figure 3 compares a simulated time-series evolution of the DW-SSFP signal using (1) EPG integrating the proposed motion operator and (2) Monte Carlo simulations (see Methods) for fixed rigid-body motion (left column) and a temporally varying motion profile characterising cardiac pulsatility (right column). Excellent agreement is found between the two approaches. Notably, motion in the EPG implementation was modelled using a single timestep per TR (Eq. [4]), with motion in the Monte Carlo implementation modelled using 100 timesteps per TR. Agreement between the two approaches suggests that the short TR of DW-SSFP enables motion to be captured as a single parameter in each TR.

**Figure 3:**
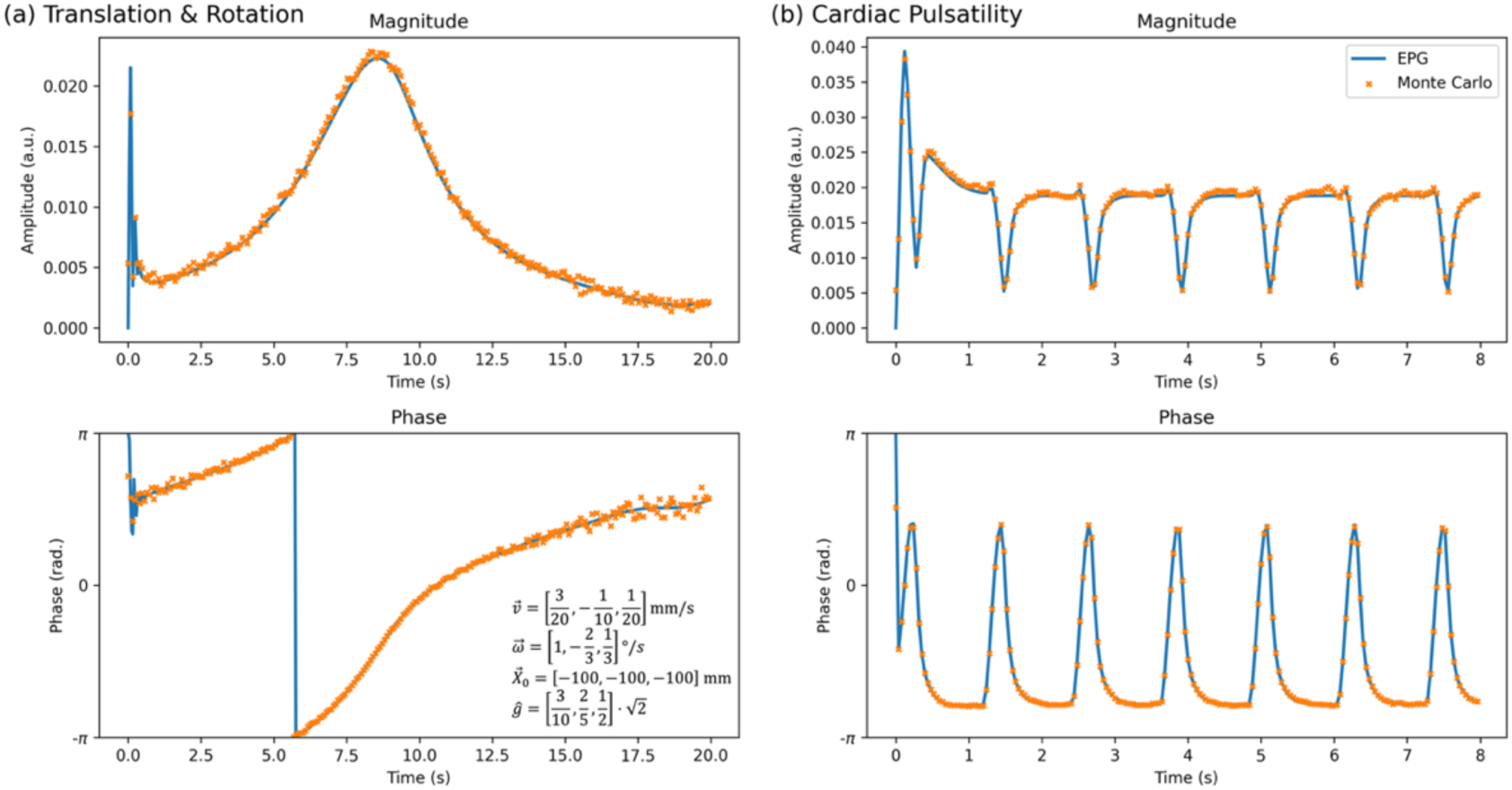
**Comparing the EPG-motion operator with Monte Carlo simulations**. Motion-corrupted DW-SSFP time-series data simulated using the proposed EPG-motion operator (blue) and Monte Carlo (orange) simulations (see Methods) for (a) fixed rigid body translation/rotation across all three dimensions and (b) cardiac pulsatility. Rigid body motion parameters associated with Eqs. [3-5] are displayed in (a). Pulsatility velocity profile (Supporting Information Figure S1) based visually on Greitz et al.^44^, simulated defining the maximum pulsatility velocity 𝑣⃗_Card_ = [0, 0, 0.4] mm/s and 𝑔< = [0,0,1], with total displacement (i.e. integral of the velocity profile) across a single cardiac cycle set equal to zero. Yellow dots correspond to the Monte-Carlo signal estimates at the end of each TR. Sequence parameters based on the experimental investigation performed in Miller & Pauly^17^, setting G = 40 mT/m, δ = 6.5 ms, α = 30°, TR = 40 ms and 𝛟 = 0°. Sample parameters were based on approximate in vivo values in brain tissue at 3T^35^, defining D = 1 ⋅ 10^−3^ mm^2^/s, T_1_ = 832 ms and T_2_ = 110 ms.

### Motion induced image corruption in DW-SSFP

To visualise the impact of motion on DW-SSFP images, we can utilise the proposed EPG-motion framework to simulate the signal evolution of individual voxels in a DW-SSFP dataset. Figure 4a-d displays synthesised DW-SSFP images assuming an instantaneous single-shot readout incorporating (a) no motion, (b) translation along a single dimension, (c) rotation along a single dimension and (d) cardiac pulsatility. Here we observe that even simple forms of rigid-body motion lead to complicated spatial changes in both signal magnitude and phase. Signal changes arising from pulsatile motion (Figure 4d) give good visual agreement to loss of signal magnitude observed previously^17,20^, with artefacts propagating into images synthesised by simulating a segmented readout (Figure 4e). Comparisons with DW-SE images acquired with the same motion profile and an instantaneous readout are displayed in Supporting Information Figure S3, where only changes in the spatial phase profile are observed.

**Figure 4:**
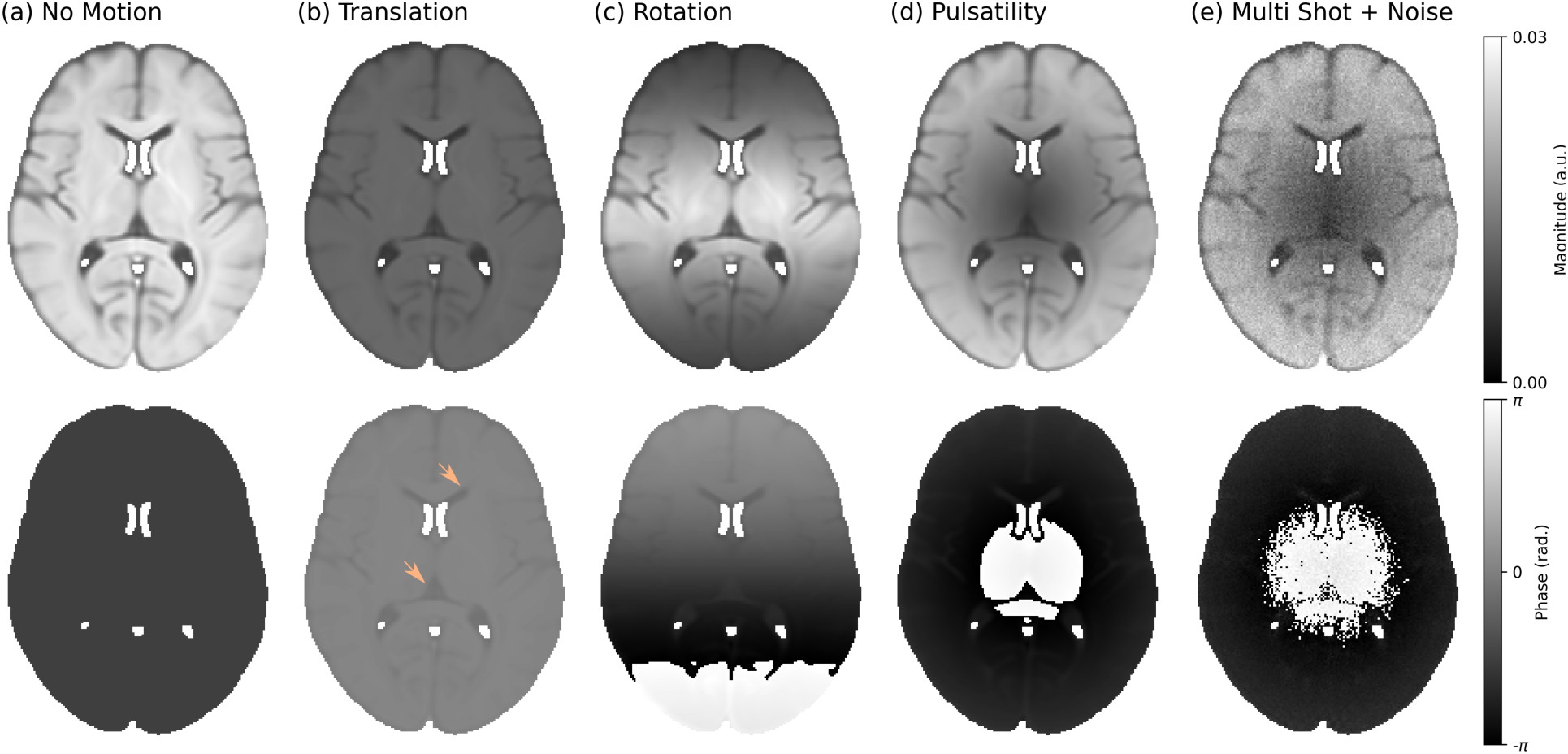
Impact of motion on DW-SSFP images. (a-d) displays simulated single-shot DW-SSFP images with an instantaneous readout incorporating (a) no motion (b) constant translation along the foot-head direction ( 𝑣⃗ = [0,0,0.2] mm/s), (c) constant rotation with the axis of rotation oriented along the left-right direction (i.e. passing through the sagittal plane) (𝜔>⃗ = [0.2,0,0] °/s), and (d) cardiac pulsatility along the foot-head direction (𝑣⃗_Card_ = [0, 0, 0.4] mm/s), all simulated with the diffusion gradient oriented along the foot-head direction (𝑔< = [0,0,1]). (e) displays a simulated DW-SSFP image acquired with a segmented readout (three-line cartesian readout per TR) incorporating cardiac pulsatility only with added Gaussian noise (SNR ∼ 20), where each readout is associated with a different component of the cardiac cycle. The light orange arrows overlaid on the phase maps in (b) display local tissue-type contrast reflecting the interplay between motion and the local diffusion coefficient that are not observed in motion-corrupted DW-SE data (Supporting Information Figure S3). Images synthesised using the HCP1065 DTI template mean diffusivity map, available as part of FSL^45^. Pulsatility timeseries profile and relative velocity map based visually on Greitz et al.^44^, provided in Supporting Information Figures S1 and S2. Sequence parameters based on the experimental investigation performed in Miller & Pauly^17^, setting G = 40 mT/m, δ = 6.5 ms, α = 30°, TR = 40 ms, 𝛟 = 0°. Relaxation parameters (set equal across all voxels) were based on approximate in vivo values in brain tissue at 3T^35^, setting T_1_ = 832 ms and T_2_ = 110 ms. Images (a-d) are displayed for the 150^th^ TR, corresponding to a simulation duration of 6 seconds. (e) was synthesised using data from the 101^st^ to the 188^th^ TR (ignoring systole), corresponding to an acquisition duration of 3.48 s. Voxels were modelled as independent (i.e. the voxel location is stationary over the time-course of the simulation).

In addition to the loss of signal magnitude in motion-corrupted DW-SSFP data, resulting phase images display local tissue-type contrast (e.g. delineation of CSF filled regions, light orange arrows in Figure 4b) that are not observed in motion-corrupted DW-SE data (i.e. the flat phase maps in Supporting Information Figure S3). This local contrast reflects the interplay between motion and the local diffusion coefficient. In the context of EPG, an increased diffusion coefficient rapidly attenuates the relative contribution of signal forming pathways associated with strong diffusion-weighting, analogous to reducing the relative contribution of signal forming pathways associated with greater motion-corruption (Supporting Information Figure S4). The local diffusion coefficient, *D*, must therefore be estimated as part of the motion-reconstruction procedure.

The DW-SSFP signal has known dependencies on local tissue relaxation properties (T^1^ & T^2^) and B^1^^32^, which must also be estimated (via complementary mapping techniques) for accurate diffusion and motion modelling. From the perspective of EPG, this property arises as T^1^, T^2^, and B^1^ values additionally modulate the relative contribution of different DW-SSFP signal-forming magnetisation pathways and interact with their relative motion sensitivity, analogous to the description of diffusion coefficients in the previous paragraph. T^1^, T^2^ and B^1^ mapping procedures using complementary imaging sequences have formed a routine part of many DW-SSFP studies estimating quantitative diffusion parameters, with a similar approach used in this study (see Methods).

### Does the DW-SSFP signal contain sufficient information to reconstruct motion-free estimates?

Earlier in this manuscript, we proposed an EPG operator for motion characterisation (Eq. [2]), validated using Monte Carlo simulations (Figure 3). Eqs. [2-4] describe motion in DW-SSFP as fully characterised provided one has knowledge of evolution of 𝑉_n_ across TRs, 𝑉(𝑡) (Eq. [4]). Importantly, whilst individual EPG pathways are associated with different levels of motion corruption in each TR, this difference solely arises from the interplay of 𝑉(𝑡) with a pathway’s *known* dephasing order, *k*. This raises the question of whether DW-SSFP data can contain sufficient information to simultaneously estimate 𝑉(𝑡) as part of the signal characterisation to reconstruct motion-corrected images. The remainder of the manuscript investigates this using both synthetic and experimental data.

## Methods

### EPG-motion Framework

Software for simulating (1) 𝑉(𝑡), and (2) the DW-SSFP signal incorporating the impact of subject motion was implemented in Python (3.13.2). The provided software allows the modelling of voxelwise and imaging timeseries DW-SSFP data based on a user-defined set of sequence parameters (*G*, 𝜏, 𝑇𝑅, 𝜃, 𝜙) and sample properties (𝑇_1_, 𝑇_2_, *D*, 𝐵_1_). When simulating 2D and 3D images (e.g. Figure 4), the software models each voxel as independent (i.e. the voxel location is stationary over the time-course of the simulation).

The software was built using NumPy (2.2.4), accelerated with the Numba compiler (0.61.2). For reference, the time required to synthesise (i.e. forward simulate) an arbitrary 𝑉(t) profile and subsequent motion corrupted DW-SSFP signal over 200 TRs is ∼5 ms (single voxel), ∼10 s (2D slice) and ∼10 min (3D volume) on a personal laptop.

### Monte Carlo Simulations

1D DW-SSFP time-series magnitude and phase signals incorporating rigid body and pulsatile motion were synthesised using Monte Carlo simulations (Figure 3). To simulate diffusion, isochromat trajectories were generated using the Camino *datasynth* function^46^, modified to produce trajectories that followed a Gaussian distribution of displacements per timestep (10^5^spins, 100 timesteps per TR). Diffusion was simulated as isotropic with a diffusion coefficient of D = 1 ⋅ 10^−3^ mm^2^/s (approximating in vivo brain tissue). To simulate motion, the location of individual isochromats was updated per timestep via translational and/or rotational operations using custom software. For the pulsatility simulations, the continuous time-series velocity profile (Supporting Information Figure S1) was averaged across the window associated with each timestep and discretised to define the isochromat displacement.

For the Monte Carlo simulations, the DW-SSFP signal arising from the isochromats was modelled using custom software, implemented in MATLAB (2023a, The MathWorks, Inc., Natick, MA).

Sequence parameters were based on the experimental investigation performed in Miller & Pauly^17^ (*G* = 40 mT/m, 𝛿 = 6.5 ms, 𝛼 = 30°, TR = 40 ms and RF phase 𝜙 = 0°), with relaxation properties based on approximate in vivo values in brain tissue at 3T^35^, T_1_ = 832 ms and T_2_ = 110 ms.

Monte Carlo simulations incorporating rigid body translational and rotational motion were synthesised defining translational velocity, 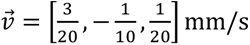, rotational velocity, 𝜔⃗ = 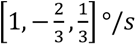, initial spatial position, 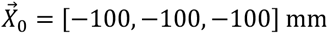, and diffusion gradient orientation, 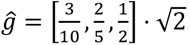. These parameters were chosen to characterise an arbitrary, constant rigid body motion across all three spatial dimensions, achieving large changes in signal magnitude and phase. The pulsatility velocity profile (characterising time-dependent motion) is visualised in Supporting Information Figure S1, setting the maximum pulsatility velocity 𝑣⃗_Card_ = [0, 0, 0.4] mm/s and *ĝ* = [0,0,1] (i.e. equivalent to the foot-head direction).

### Parameter Estimation from Simulated DW-SSFP Data (Monte Carlo)

Using the proposed EPG-motion framework, the diffusion coefficient (*D*), phase offset (𝜙), and 𝑉(t) profile were estimated from 1D DW-SSFP timeseries data arising from the Monte Carlo simulations incorporating pulsatility (Figure 3b). The phase offset (𝜙) reflects deviations in the true phase offset versus the defined RF phase angle, equivalent in simulations (𝜙 = 0°) but differing experimentally (arising from spatial B^0^ and B^1^ profiles). To more accurately reflect experimental conditions, Gaussian noise was added to the simulated signal, leading to a final SNR of 20.

The fitting was implemented in Python (3.13.2) using the Trust Region Reflective algorithm (SciPy *curve_fit* 1.15.2). Specifically, the EPG-motion framework was incorporated as a forward model, with the parameter estimation procedure identifying parameters that minimised the cost function with the motion-corrupted Monte-Carlo DW-SSFP signal. To accelerate the fitting procedure, an analytical Jacobian function was derived for all parameters (except for the derivative with respect to *D*, which was estimated numerically). Initialisation parameters and fitting bounds are provided in Supporting Information Table S1, with no additional constraints or regularisation incorporated.

The simulated DW-SSFP timeseries data consisted of 200 TRs. Parameter estimation was performed using a time window of 100 TRs of steady-state signal (after 100 TRs, from 3.96 s in Figure 3b). A single motion parameter was estimated per TR, corresponding to 102 parameters in total (*D* + 𝜙 + 𝑉(t)). To facilitate estimation of 𝑉(t) the parameter estimation region was split into two components corresponding to (1) dummy measurements (first 25 TRs) and (2) measured signal (remaining 75 TRs). The dummy measurements reflect a series of TRs which were not incorporated into the cost function evaluation. This is important as motion effects build up over several TRs in DW-SSFP. Specifically, the dummy region makes it possible to accurately characterise the motion-corrupted signal from the first TR of the measured signal region by allowing the signal to reach a steady-state incorporating the effects of motion prior to this timepoint. Ten repeats were performed with different noise instances to characterise the robustness of the fitting approach. The fitting procedure averaged at ∼4 seconds per noise instance (i.e. per 1D timeseries) on a personal laptop (Macbook Pro, Sonoma 14.5, M1, 16GB RAM).

To understand the impact of incorporating motion into the parameter estimation procedure, an identical fitting approach was implemented without estimation of the 𝑉(t) operator (i.e. conventional EPG). This corresponds to the estimation of two parameters per simulation (*D* + 𝜙).

### Estimation of *D* as a Function of SNR and Pulsatile Velocity

To assess the robustness of characterising *D* from motion-corrupted DW-SSFP timeseries data, simulations were performed incorporating pulsatile motion for a range of SNR levels (10, 20, 50, + no noise) and maximum pulsatility velocities (0 to 1.5 mm/s in 0.3 mm/s increments), where 0 mm/s corresponds to a no motion case and 1.5 mm/s corresponds to a previous experimental estimate of maximum pulsatile velocity in brain tissue^44^. Ten repeats were performed per SNR level and velocity instance (240 simulations total).

### Experimental Data Acquisition

2D time-series DW-SSFP data were acquired in the brain in three healthy volunteers on a 3T Siemens Prisma scanner (32 channel head coil). Data was acquired in a single axial slice for Subject 1, with axial and coronal slices acquired in Subjects 2 and 3. The subjects gave informed consent with data acquisition in accordance with local ethics committees. Sequence parameters were optimised for a target effective b-value, b_eff_ = 500 s/mm^2^, setting *G* = 63.6 mT/m, 𝛿 = 3.08 ms, TR = 26 ms, 𝛼 = 31°, 𝜙 = 0°, TE = 15 ms, BW = 2405 Hz/Pix, resolution = 6 × 6 × 6 mm^3^, matrix size = 32 × 32 with a single-shot Cartesian EPI readout (16 ms duration). b^0^-equivalent data were also acquired with *G* = 26.7 mT/m, producing images that achieve the DW-SSFP spoiling condition with low diffusion weighting (b_eff_ = 100 s/mm^2^). These are henceforth referred to as b^100^, with the b_eff_ = 500 s/mm^2^ data referred to as b^500^.

To ensure the signal reached a steady-state, dummy measurements were performed for 5 seconds prior to data acquisition. Sequence parameters were chosen via an optimisation routine that maximised SNR-efficiency for the target effective b-value^47^. To evaluate the relationship between the DW-SSFP timeseries data and cardiac pulsatility, pulse oximeter recordings were also acquired.

The time-series data consisted of 384 consecutive measurements in a single axial slice of the brain (10 second acquisition). A fully sampled image was acquired per TR (i.e. equivalent to a ‘single-shot’ readout) to characterise the temporal dynamics of the DW-SSFP signal independent of a segmented readout scheme. 12 diffusion directions were acquired, with both b^100^ & b^500^ volumes acquired per direction. The raw data (.*dat*) was exported from the scanner and reconstructed using *mapVBVD*^48^, with coil-combination performed using SVD and ESPIRiT^49,50^

The DW-SSFP signal has known dependencies on tissue relaxation (T^1^ and T^2^) properties and the B^1^ profile. Complementary T^1^, T^2^, and B^1^ maps were estimated in the same subject using a spin echo echo planar imaging (EPI) sequence with different inversion preparations (ep2d_se), multi-echo spin-echo (se_mc), and 3DREAM^51^. Full details of the sequence parameters and processing are provided in Supporting Information Table S1.

To facilitate comparisons, 2D DW-SE EPI data was additionally acquired within the same subjects, closely matching the acquisition scheme of the DW-SSFP data (b_0_and 𝑏 = 500 s/mm^2^, 12 diffusion directions). Full details of the DW-SE EPI sequence parameters and image processing are provided in Supporting Information Table S1.

### Parameter Estimation from Experimental DW-SSFP Data

The diffusion coefficient (*D*), signal amplitude (𝑆_0_), phase offset (𝜙), and 𝑉(t) operator were estimated from experimental DW-SSFP timeseries data using the proposed EPG-motion framework. The fitting procedure was implemented in Python (3.13.2) as described above, fitting simultaneously to the time-series b^100^ and b^500^ data. Initialisation parameters and fitting bounds are provided in Supporting Information Table S3, with no additional constraints or regularisation incorporated. Fitting was performed independently for each voxel and diffusion direction, estimating a single diffusion coefficient per voxel.

For the fitting procedure, DW-SSFP timeseries data was simulated for 509 TRs. This consisted of 100 TRs to ensure the simulated signal reached a steady-state, 25 TRs dummy measurements, and 384 TRs where comparisons were made to the acquired experimental data. A single motion parameter was estimated per TR, corresponding to the estimation of 822 parameters per voxel (*D* + 𝑆_0_ + 𝜙_𝑏100_+ 𝜙_𝑏500_+ 𝑉_𝑏100_(t) + 𝑉_𝑏500_(t)). Parameter estimation required ∼30 seconds per voxel on a personal laptop (Macbook Pro, Sonoma 14.5, M1, 16GB RAM). Finally, a tensor was reconstructed from the estimated *D* maps acquired across all 12 orientations using custom software.

To understand the impact of incorporating motion into the parameter estimation procedure, an identical fitting approach was implemented without estimation of the 𝑉(t) operator (i.e. conventional EPG), estimating four parameters per simulation (*D* + 𝑆_0_ + 𝜙_𝑏100_+ 𝜙_𝑏500_).

## Results

Figure 5 displays the estimated DW-SSFP time-series and 𝑉(t) operator using the proposed EPG-motion framework, arising from Monte Carlo simulations of the steady-state DW-SSFP signal incorporating pulsatile motion with added Gaussian noise. Excellent agreement is found over the fitted data region (green box), indicating that simultaneous estimation of 𝑉(𝑡) and D is possible from DW-SSFP timeseries data.

**Figure 5:**
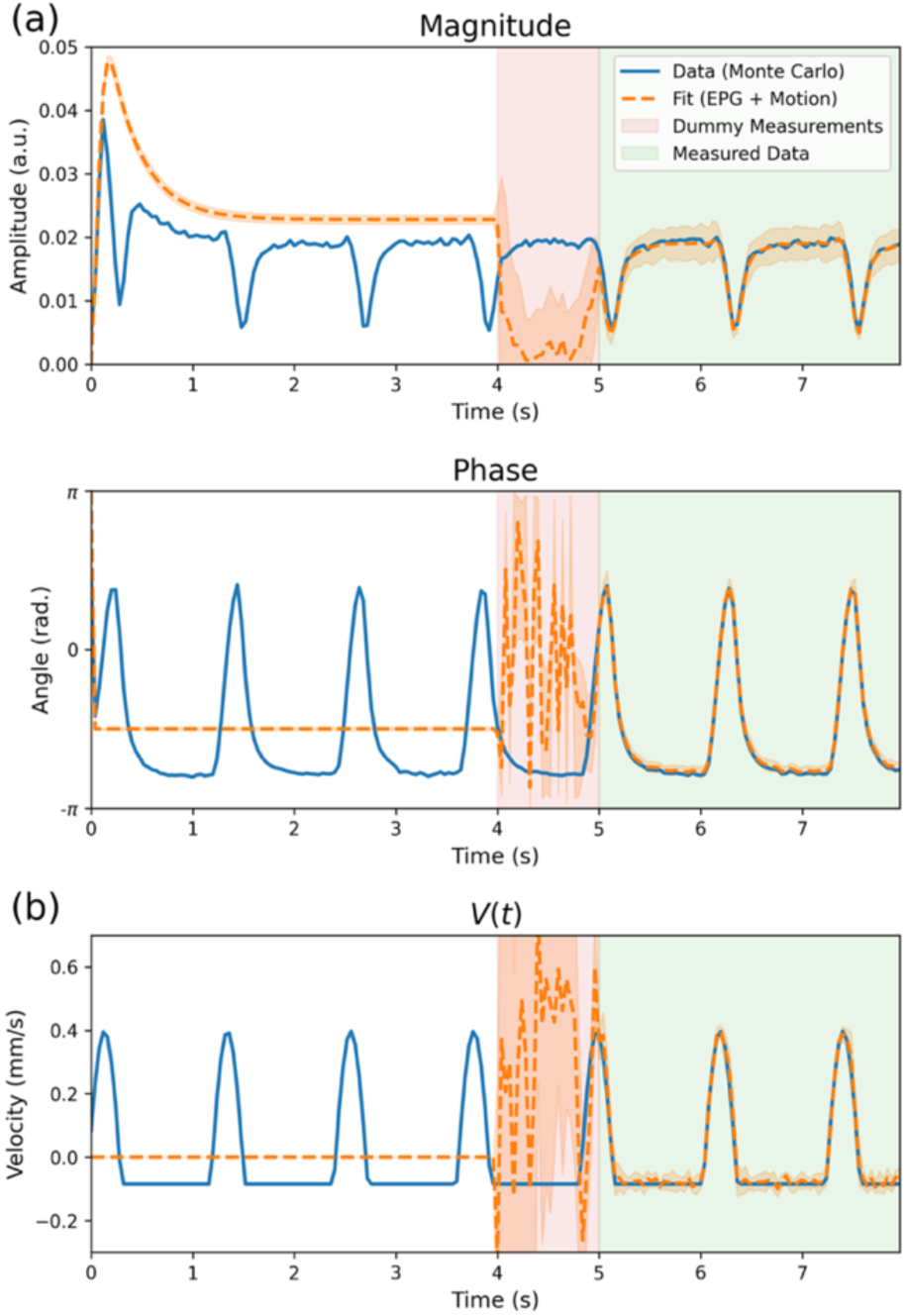
Estimation of 𝑽(𝒕) from simulated DW-SSFP time-series data. (a) displays DW-SSFP time-series magnitude and phase data (blue) incorporating cardiac pulsatility + Gaussian noise (SNR = 20) synthesised using Monte Carlo simulations. Fitting to the data via the proposed EPG-Motion framework accurately characterises the DW-SSFP time-series profile (orange) in the measured data region (green box). (b) The estimated 𝑉(𝑡) time-series profile (reflecting the component of instantaneous velocity in the direction of the diffusion gradient across TRs) closely matches the ground truth in the measured data region (green), reflecting the possibility of characterising motion solely from the DW-SSFP signal. Dummy measurements (pink box) reflect a series of TRs where no comparison to the simulated data was performed. The white box reflects a series of TRs where the simulation reached a steady state. Here the dashed orange line represents the average estimate over the 10 noise repeats, with the orange shaded region surrounding the fitted line corresponding to the standard deviation over the 10 noise repeats.

When considering diffusion MRI acquisitions, the key parameter we want to accurately characterise is the diffusion coefficient, D. By means of comparison, given the simulated coefficient (D = 1 ⋅ 10^−3^ mm^2^/s), fitting to the simulated data with no estimation of 𝑉(𝑡) (i.e. conventional EPG) yielded D = (2.00 ± 0.02) ⋅ 10^−3^ mm^2^/s. Fitting with the proposed EPG-motion framework estimated D = (0.99 ± 0.04) ⋅ 10^−3^ mm^2^/s.

Data acquisition for a DW-SSFP experiment does not begin at the first TR, with motion during the build-up to a steady state leading to motion corruption of the DW-SSFP signal from the first TR of data acquisition. This is visible in the measured data region of Figure 5 (green box), where the signal amplitude in the first TR is below the amplitude predicted of a motion-free DW-SSFP signal (Figure 5 – dashed orange line). If the estimation of motion only begun at the first TR of the measured data region (green box), it would not be possible to accurately estimate the motion profile or the signal in the first few TRs of the measured data region, as it would have to immediately transition from a motion-free steady-state to a measurement representing the evolution of a motion-corrupted DW-SSFP signal that has arisen over several TRs.

To account for this, the parameter estimation routine allows for motion to be incorporated into the TRs prior to the measurement region, here referred to as dummy measurements (pink box). In this box no cost function evaluations are made, with the presence of dummy measurements enabling the signal to be accurately represented from the first TR of the measured data region (green box). The choice of 25 TRs corresponds to a single second of data, reflecting the approximate time required to return to steady state for an instantaneous motion operation (Figure 2) based on the simulated parameters investigated here. The arbitrary evolution of the signal/motion profiles in the dummy measurement region (pink box) arises as no cost-function evaluations are made, meaning parameter estimation is poorly conditioned in this region. However, the signal/motion profiles begin to more closely represent the ground truth as the measured data region is approached (right hand side of the pink box), with the accurate representation of the motion profile from the first TR of the measured data region (green) demonstrating that their inclusion is working as intended.

Figure 6 displays the estimated diffusion coefficient from simulated pulsatile motion with varying SNR levels and maximum pulsatility velocity. Here we identify that the incorporation of motion properties in parameter estimation substantially reduces the misestimation of D. Specifically, no characterisation of subject motion (Figure 6a) led to biases in D of up to 275% across the investigated velocity/SNR regime, increasing as a function of the maximum pulsatile velocity. An equivalent investigation incorporating the estimation of motion parameters using the EPG-motion framework (Figure 6b) led to a maximum bias of 12%.

**Figure 6:**
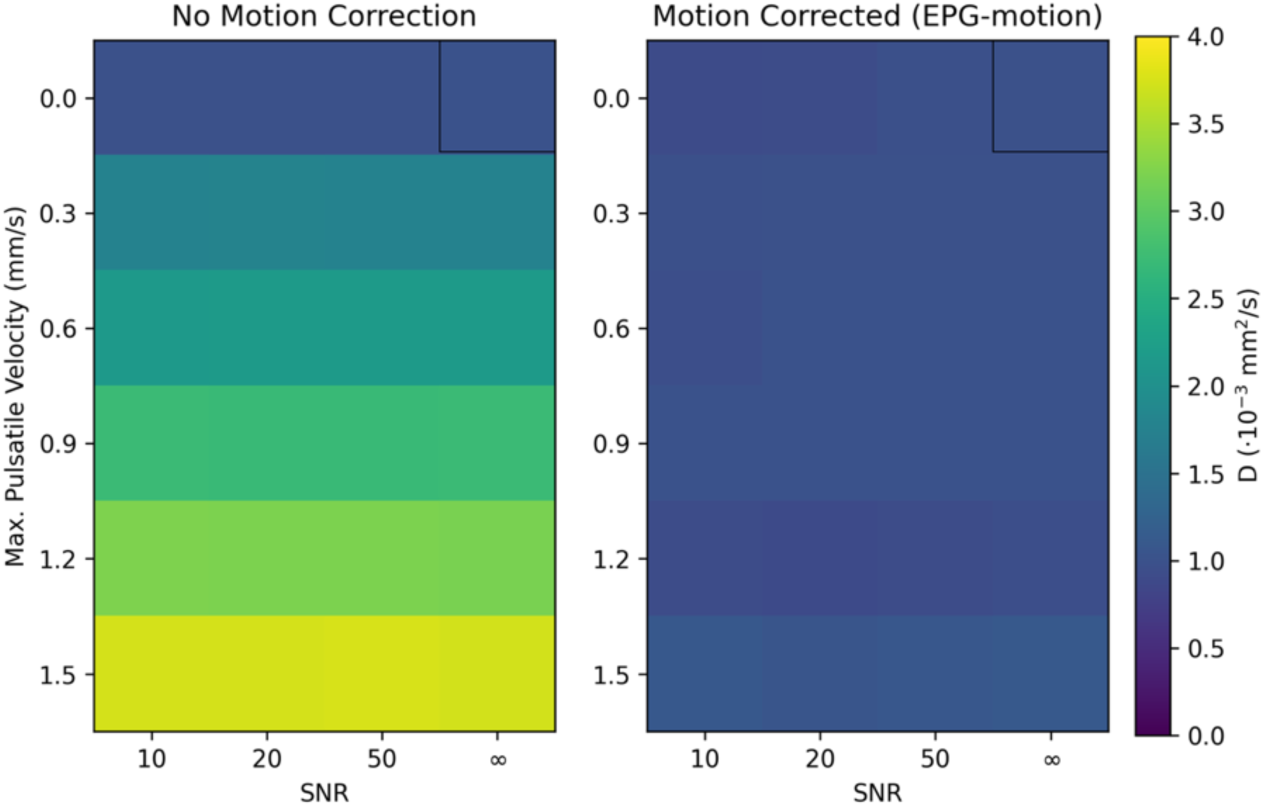
Characterisation of D as a function of SNR and maximum pulsatile velocity amplitude. (a) and (b) display the estimated diffusion coefficient, D, derived from DW-SSFP timeseries data as a function of SNR (x-axis) and maximum pulsatile velocity (y-axis). The two plots correspond to parameter estimation assuming (a) no motion correction and (b) motion correction using the proposed EPG-motion framework. The black box in the top right corresponds to the ground truth estimate (no motion, SNR = ∞). No incorporation of motion in parameter estimation (a) led to a maximum bias of 275% in the estimation of D across the investigated velocity/SNR regime, increasing as a function of the maximum pulsatile velocity. Simultaneous estimation of motion parameters (b) led to a maximum bias of 12%. Figure displays the average D calculated from 10 simulated DW-SSFP timeseries repeats per SNR/velocity level.

The motion-free simulations (Figure 6 – top row) with no added noise (Figure 6 – black box) estimated 0.998 ⋅ 10^−3^ mm^2^/s (with motion correction) and 1.000 ⋅ 10^−3^ mm^2^/s (no motion correction). A small bias was observed in the diffusion coefficient estimation at reduced SNR levels when incorporating motion correction (∼8% at SNR = 10, Figure 6 - top left) for the motion-free simulations, which were not present in the parameter estimation procedure without motion correction (∼0.4% at SNR = 10). This reflects an increased sensitivity of the proposed EPG-motion framework to data noise.

Figure 7 displays an example DW-SSFP timeseries and 𝑉(𝑡) profile estimated in a single voxel of experimental data acquired in the human brain. The timeseries corresponds to the voxel in the thalamus, with the diffusion gradient oriented along the anterior-posterior axis. The experimental timeseries data displays consistent characteristics to the DW-SSFP simulations incorporating cardiac pulsatility, with the periodicity of the DW-SSFP signal and 𝑉(𝑡) profile consistent with the recorded locations of pulse oximeter triggers.

**Figure 7:**
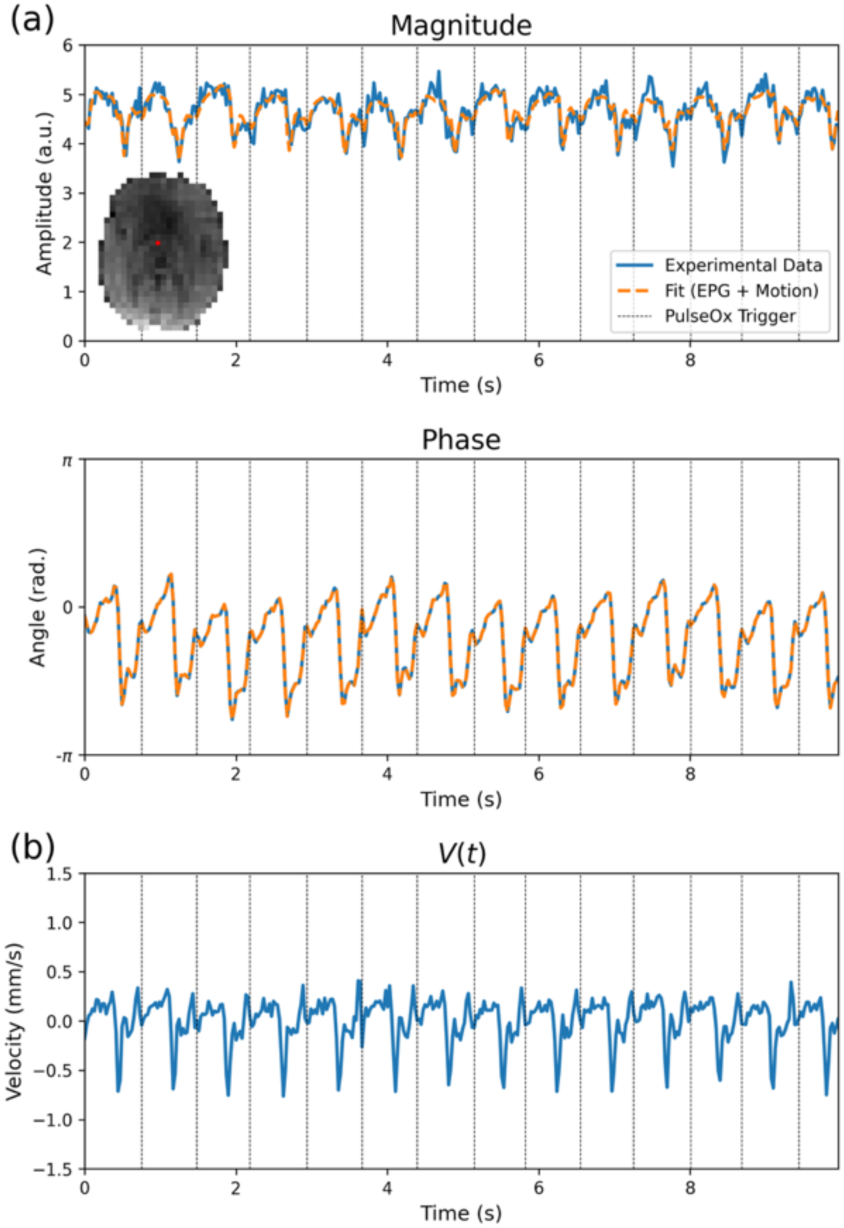
DW-SSFP signal and 𝑽(𝒕) profile in a single voxel of experimental 𝐛_𝐞𝐟𝐟_ = 𝟓𝟎𝟎 𝐬/𝐦𝐦^𝟐^data. The experimental DW-SSFP signal profile (a) (blue lines) displays periodicity consistent with the timings of pulse oximeter triggers (black vertical lines) acquired as part of the acquisition. Fitting to the experimental data using the proposed EPG-motion framework (orange line) gives excellent agreement to the experimental data, with the estimated velocity profile (b) displaying consistent periodic changes in tissue velocity, consistent with pulse oximeter trigger locations. Displayed timeseries corresponds to a single voxel in the thalamus (visualised in inset figure of experimental data in (a) – red voxel) from Subject 1, with the diffusion gradient oriented along the anterior-posterior axis. 𝑉(t) defined as the component of instantaneous velocity in the direction of the diffusion gradient across TRs (Eq. [4]).

Figure 8 displays the DW-SSFP timeseries magnitude data (b^500^) across a portion of the cardiac cycle for Subject 1 with the diffusion gradient oriented along the (a) left-right, (b) anterior-posterior and (c) foot-head direction. Subtle changes in signal amplitude are visible across the timeseries in (a) and (b), with considerable changes in signal amplitude along the foot-head direction (c). Here, even a modest b-value of 500 s/mm^2^ leads to a substantial loss of signal along the foot-head direction in central and anterior brain regions.

**Figure 8:**
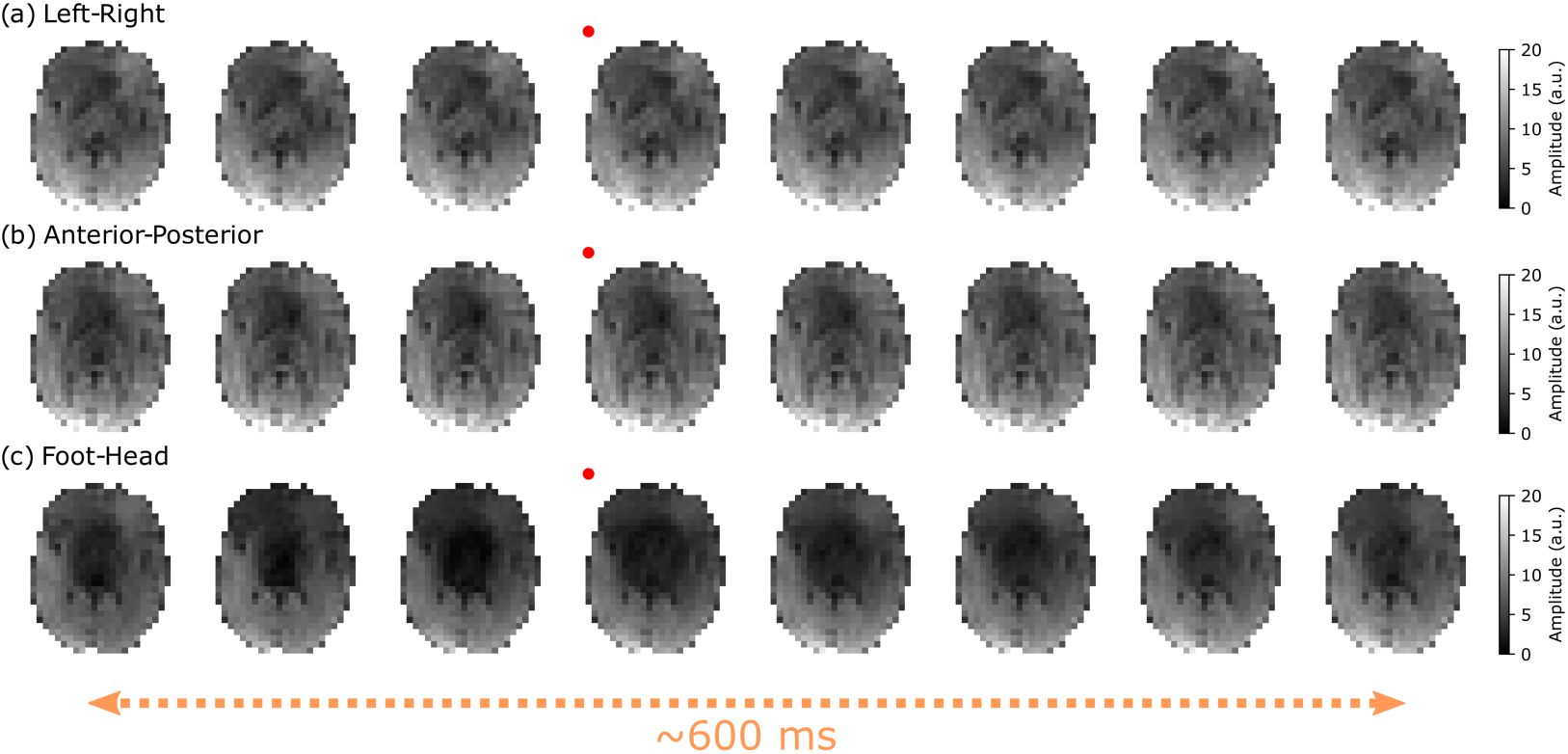
Experimental DW-SSFP timeseries images (Subject 1). (a-c) correspond to example b500 timeseries data acquired with left-right (a), anterior posterior (b) and foot-head (c) gradient orientations in Subject 1. The change in signal is most prominent along the foot-head orientation, characterised by a substantial loss of signal in central and anterior brain regions. Here the red dots indicate the image acquired at the timepoint of the pulse oximeter trigger, with the x-axis reflecting a subset of data acquired over 624 ms, with a gap of 78 ms between images (three TRs). Corresponding figures for Subjects 2 and 3 are provided in Supporting Information Figures S5 and S6.

Equivalent figures are provided in Supporting Information for Subject 2 (Figure S5) and Subject 3 (Figure S6) for axial and coronal acquisitions. Coronal acquisitions display considerable motion-induced signal loss along the foot-head direction in central brain and brain stem regions. All three subjects offer similar orientation-dependent contrast, with Subject 2 displaying reduced signal levels in the anterior brain in comparison to Subject 1, and Subject 3 demonstrating a much greater loss of signal along the foot-head direction for both axial and coronal acquisitions, indicative of increased subject motion.

Figure 9 displays the corresponding 𝑉(𝑡) maps for Subject 1 estimated from the same portion of the cardiac cycle with the diffusion gradient oriented along the (a) left-right, (b) anterior-posterior and (c) foot-head direction. Here we observe smooth changes (both temporally and spatially) in 𝑉(𝑡).

**Figure 9:**
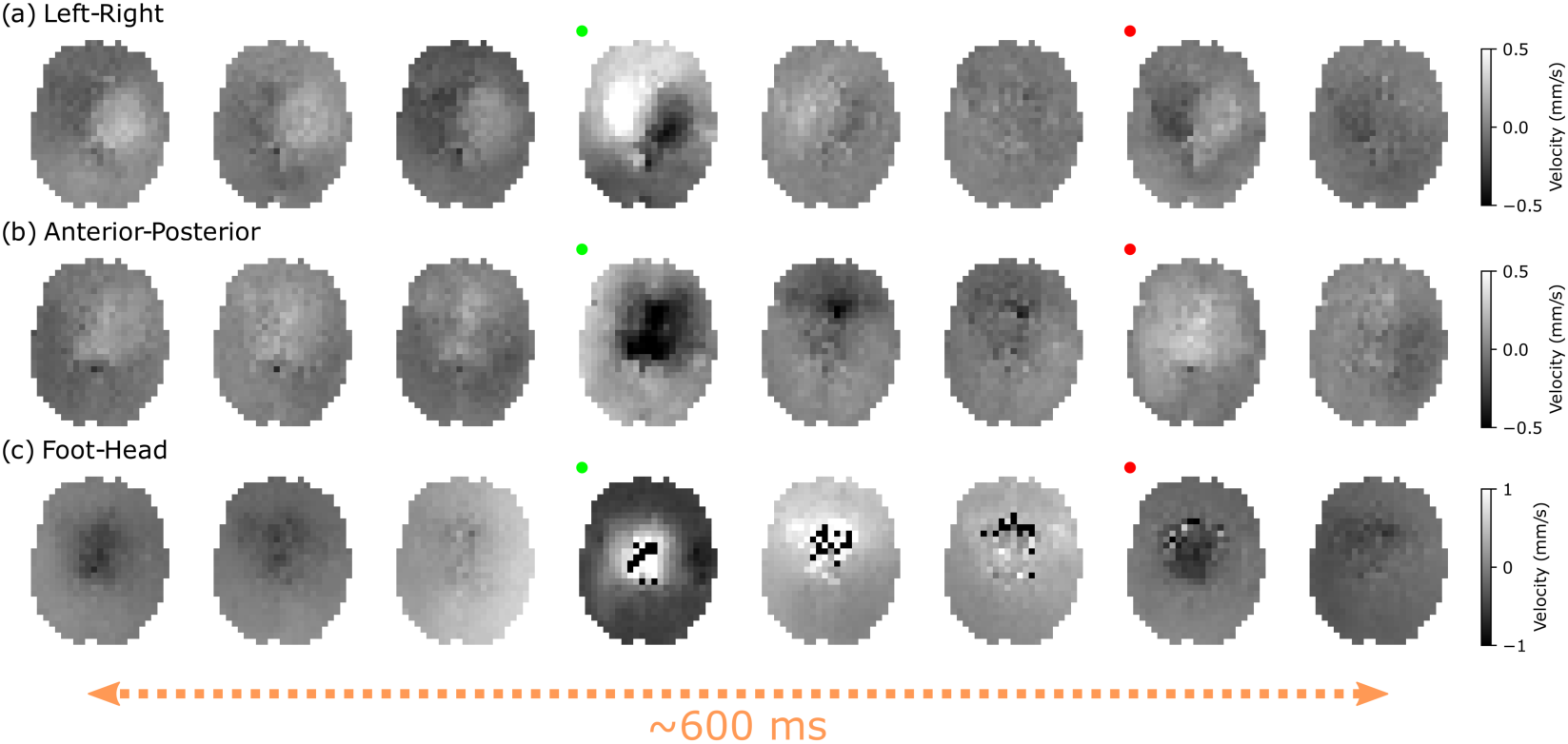
Estimated 𝑽(𝒕) maps (Subject 1). (a-c) correspond to example 𝑉(𝑡) spatial maps estimated from the experimental b500 DW-SSFP data acquired along left-right (a), anterior-posterior (b) and foot-head (c) gradient orientations in Subject 1. Here the red dots indicate the image acquired at the timepoint of the pulse oximeter trigger, with the green dots indicating peak brain tissue velocity changes. The x-axis reflects a subset of data acquired over 624 ms, with a gap of 78 ms between images (three TRs). 𝑉(t) defined as the component of instantaneous velocity in the direction of the diffusion gradient across TRs (Eq. [4]). Black/white voxels within the brain correspond to velocities exceeding the colour bar limits (right hand side). Corresponding figures for Subjects 2 and 3 are provided in Supporting Information Figures S7 and S8.

Importantly, each voxel in these images has been processed independently, with the arising spatial smoothness reflective of the consistency of velocity estimation across individual voxels. A notable exception arises from voxels in the centre of the brain along the foot-head direction (Figure 8c) where motion parameters could not be reliably estimated during peak systole (see Discussion).

The 𝑉(𝑡) maps display a consistent pattern of sharp velocity increases (associated with systole) followed by reduced velocities during diastole. The asymmetric 𝑉(𝑡) profile associated with the left-right gradient orientation is consistent with previous observations^52,53^, and an increased absolute velocity in the central brain along the anterior-posterior gradient orientation is consistent with Greitz et al^44^.

Supporting Information Figures S7 and S8 provide equivalent spatial 𝑉(𝑡) maps for Subjects 2 and 3. Coronal 𝑉(𝑡) profiles are consistent with the axial 𝑉(𝑡) profiles, with an asymmetric 𝑉(𝑡) profile for the left-right gradient orientation, and increased absolute velocity in the central brain for the anterior-posterior gradient orientation. Subjects 2 and 3 provide smooth 𝑉(𝑡) maps for left-right and anterior-posterior gradient orientations, with noisier 𝑉(𝑡) estimates in comparison to Subject 1 along the foot-head orientation where parameters could not be reliably estimated. These findings are consistent with areas of reduced signal in the corresponding data (Supporting Information Figures S5 and S6).

Figure 10 displays tensor estimates reconstructed from the 12 diffusion gradient orientations for (a) DW-SSFP timeseries data without motion correction, (b) DW-SSFP timeseries data with motion correction, and (c) DW-SE EPI data acquired in Subject 1. Without performing motion correction, it is not possible to reconstruct informative tensor estimates. Correction using the proposed EPG-motion framework leads to consistent tensor estimates across the brain (Figure 10b), with excellent agreement to DW-SE EPI data (Figure 10c)Finally, Figures 11 and 12 display tensor estimates reconstructed from Subjects 2 and 3 for both axial and coronal slices. The performance of the reconstruction from Subject 2 is consistent with Subject 1, with some small residual spatial biases in the centre of the brain where motion is predicted to be greatest (Supporting Information Figure S7). Subject 3 displays considerable biases prior to motion correction (Figure 12 top rows), with L1 estimates exceeding the presented data range (5 ⋅ 10^−3^ mm^2^/s). Following motion correction (middle rows), diffusivity estimates are more in line with the DW-SE EPI estimates (bottom rows). However, residual spatial biases are present across the slice, most apparent in the 𝑉^<⃗^_1_ maps.

**Figure 10:**
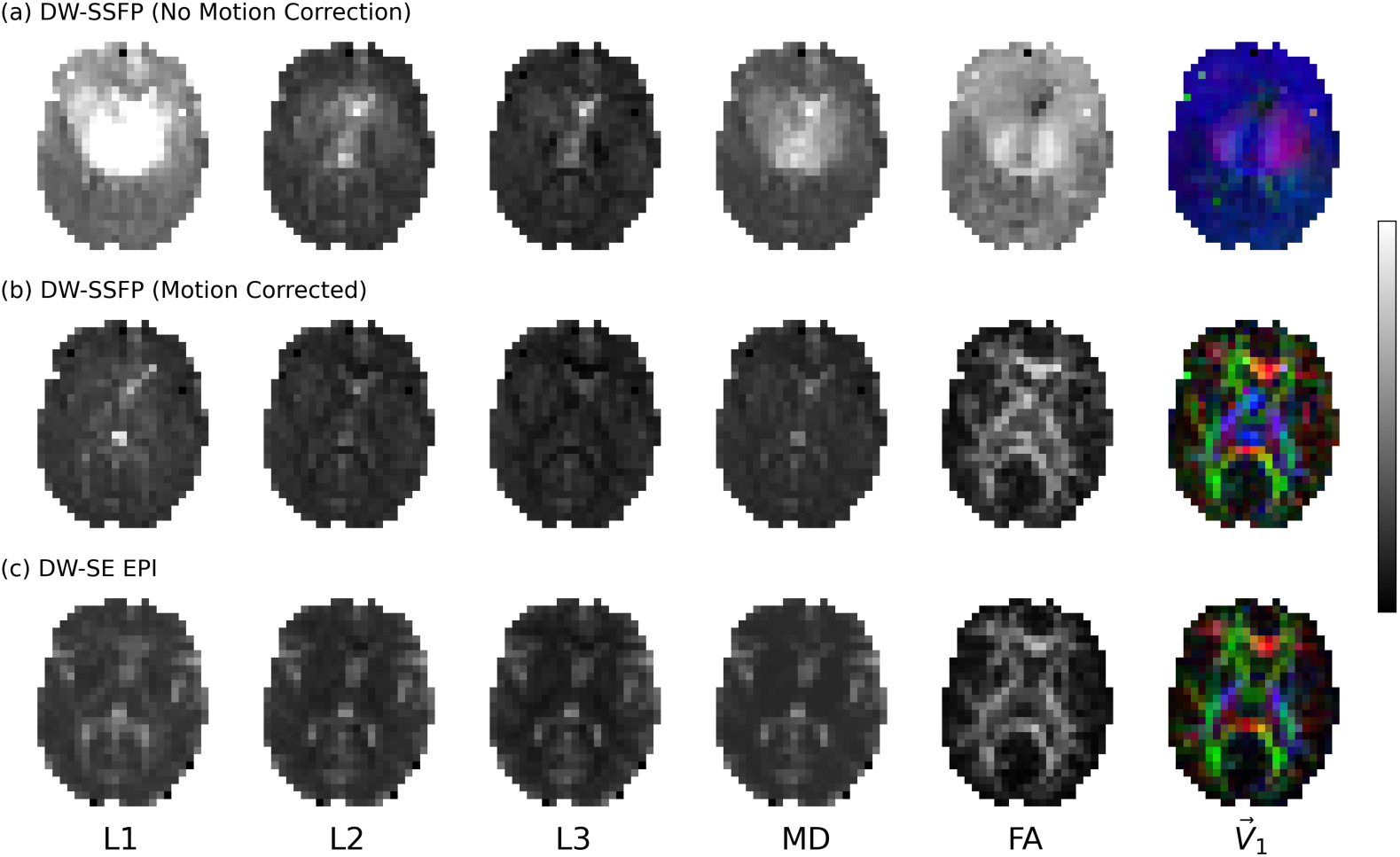
Tensor Outputs (Subject 1). Comparison of tensor outputs for (a) DW-SSFP data without motion correction, (b) DW-SSFP data with motion-correction (via the EPG-motion framework) and (c) comparison DW-SE EPI data in Subject 1. Without motion-correction (a), DW-SSFP tensor estimates display large spatial biases. Motion-corrected DW-SSFP data (b) display excellent visual agreement with complementary DW-SE EPI estimates (c), attenuating spatial biases. Tensors reconstructed from individual diffusion coefficient maps estimated per orientation using custom code. Blue = Foot-Head, Red = Left-Right, Green = Anterior-Posterior. Greyscale colour bar (right) reflects range between 0 to 5 ⋅ 10^−3^ mm^2^/s (L1/L2/L3/MD) and 0 to 1 (FA). Corresponding figures for Subjects 2 and 3 are provided in Figures 11 and 12.

**Figure 11:**
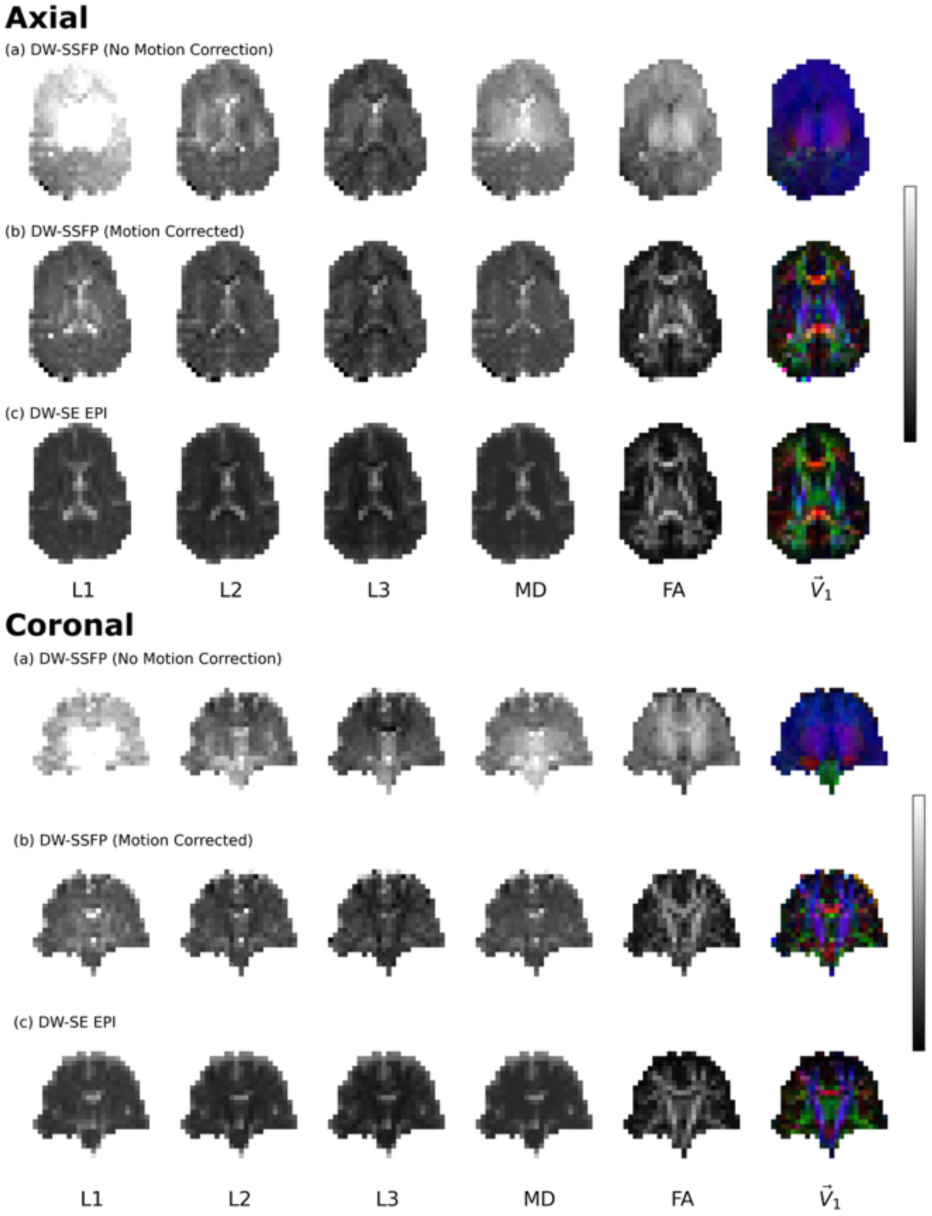
Tensor Outputs (Subject 2). Equivalent to Figure 10, here we display tensor outputs for (a) DW-SSFP data without motion correction, (b) DW-SSFP data with motion-correction (via the EPG-motion framework) and (c) comparison DW-SE EPI data for an axial (top) and coronal (bottom) slice in Subject 2. Here we observe similar motion correction performance to Subject 1, with a small residual spatial inhomogeneity remaining in the centre of the brain, most apparent in the 𝑉>⃗_1_ maps. Tensors reconstructed from individual diffusion coefficient maps estimated per orientation using custom code. Blue = Foot-Head, Red = Left-Right, Green = Anterior-Posterior. Greyscale colour bar (right) reflects range between 0 to 5 ⋅ 10^−3^ mm^2^/s (L1/L2/L3/MD) and 0 to 1 (FA), equivalent to the ranges in Figure 10.

**Figure 12:**
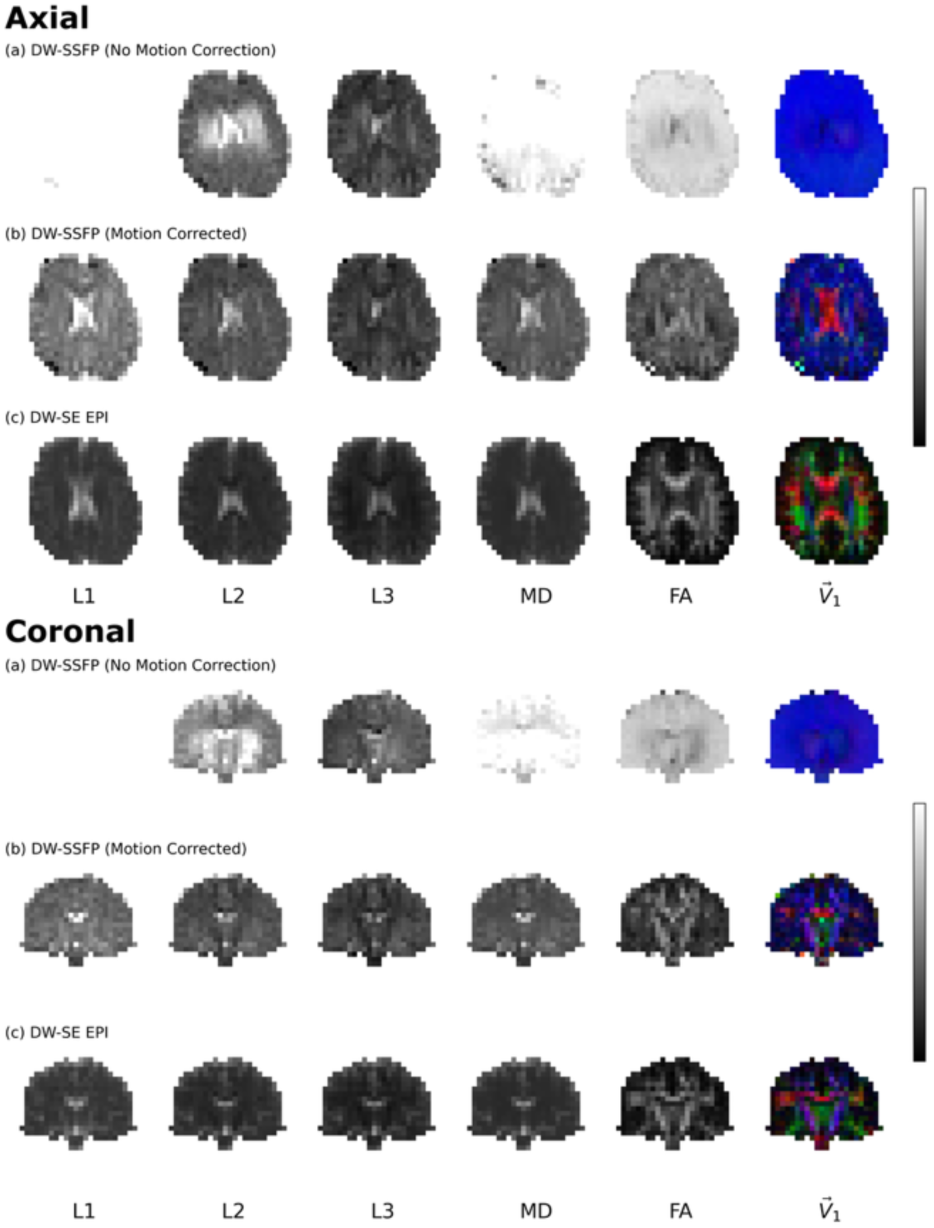
Tensor Outputs (Subject 3). Equivalent to Figures 10 and 11, here we display tensor outputs for (a) DW-SSFP data without motion correction, (b) DW-SSFP data with motion-correction (via the EPG-motion framework) and (c) comparison DW-SE EPI data for an axial (top) and coronal (bottom) slice in Subject 3. Here we observe poorer motion correction performance in comparison to Subjects 1 and 2, with greater residual spatial inhomogeneities remaining in the axial and coronal slices, most apparent in the 𝑉>⃗_1_ colour maps. Tensors reconstructed from individual diffusion coefficient maps estimated per orientation using custom code. To account for spurious diffusion coefficient estimates, the tensor reconstruction of the axial DW-SSFP data only was inversely weighted by the variance the diffusion coefficient estimate (provided by SciPy *curve_fit*). Blue = Foot-Head, Red = Left-Right, Green = Anterior-Posterior. Greyscale colour bar (right) reflects range between 0 to 5 ⋅ 10^−3^ mm^2^/s (L1/L2/L3/MD) and 0 to 1 (FA) equivalent to the ranges in Figures 10 and 11.

## Discussion

### EPG-motion Framework

The proposed EPG-motion framework demonstrates that the impact of rigid body and pulsatile motion on the underlying magnetisation distribution of DW-SSFP leads to substantial changes in DW-SSFP signal magnitude and phase (Figures 3 and 4). Acquired experimental data incorporating a single-shot readout agrees visually with forward simulations arising from the EPG-motion framework, indicative of motion-induced signal corruption arising from magnetisation distribution of DW-SSFP (Figures 7-9). Without correction incorporating the proposed EPG-motion framework, diffusion coefficients are overestimated both theoretically (Figure 6a) and experimentally (Figures 10a-12a), with the degree of misestimation additionally dependent on the diffusion gradient orientation (Figures 10a to 12a).

The proposed framework demonstrates that it’s possible to model and correct for the impact of subject motion arising from the underlying magnetisation distribution retrospectively based on timeseries data capturing the evolution of the DW-SSFP signal, yielding parameter estimates with more consistent spatial contrast (Figures 10b to 12b) and agreement with complementary diffusion imaging methods (Figures 10c to 12c). Motion-corrected data using the proposed EPG-motion framework attenuates the blue bias visible in earlier efforts to estimate tensors from motion-corrected DW-SSFP data^20^, arising from the predominant motion-sensitivity of the DW-SSFP signal along the foot-head direction^20,21^.

The proposed motion operator introduced in the Theory section (building on a previous description^34^) is not specific to DW-SSFP and can be incorporated into EPG signal representations to investigate the impact of subject motion on arbitrary MRI sequences. Other methods that are anticipated to benefit from the concepts introduced in this manuscript include DW-DESS^39–41^ and MR fingerprinting^42,43^ methods, which are similarly sensitive to motion-induced signal loss arising from the weighted sum of magnetisation components with different evolution histories.

### Motion Characterisation during Systole

The estimated 𝑉(t) maps (Figure 9 and Supporting Information Figures S7 and S8) display smooth and consistent contrast across the brain during diastole and systole along left-right and anterior-posterior gradient orientations. 𝑉(t) estimates along the foot-head gradient orientation (where subject motion is predicted to be largest) were noisier, with visualisation of the DW-SSFP data (Figure 8c and Supporting Information Figures S5c and S6c) identifying that noisy motion parameter estimates are also associated with regions of low SNR. This suggests that parameter misestimation during systole is a likely consequence of (1) noise and/or (2) parameter degeneracies associated with rapid signal changes in high-velocity tissue regions.

To investigate this, Supporting Information Figures S9-11 displays the spatial 𝑉(t) maps estimated from the experimental b^100^ (b_eff_ = 100 s/mm^2^) data for Subjects 1-3. The spatial 𝑉(t) maps display consistent contrast with the b^500^ (b_eff_ = 500 s/mm^2^) 𝑉(t) maps (Figure 9), without noisy velocity estimates observed when the diffusion gradient is oriented along the foot-head axis. Figure S12 displays an example DW-SSFP timeseries and 𝑉(𝑡) profile from the b^100^ dataset in a central brain (thalamus) voxel in Subject 1, demonstrating a periodic profile consistent with the pulse oximeter estimates.

The b^500^ data have intrinsically lower SNR and larger motion-induced signal changes versus the b^100^ data. Figure S13 displays the DW-SSFP timeseries and 𝑉(𝑡) profile for b^500^ data equivalent to Figure S12. Here we observe an excellent fit, with rapid magnitude/phase changes in the DW-SSFP signal and large non-physical oscillatory velocity estimates during systole exceeding 3 mm/s. Whilst the signal-forming mechanisms of DW-SSFP create a nontrivial relationship between sequence parameters and the maximum velocity that can be reliably estimated, the excellent fit combined with a consistent, oscillatory velocity profile during systole is indicative of degeneracies in the relationship between 𝑉(t) and the measured signal.

For Subjects 1 and 2, misestimation of motion parameters during peak systole along the foot-head gradient orientation does not appear to significantly impact estimation of *D* both experimentally (Figures 10 and 11) and in simulation (Figure 6 and Supporting Information Figure S14). However, for Subject 3 spatial biases are still present following motion correction. To investigate this further, Figure 13 displays the average timeseries velocity for all three subjects along the foot-head gradient orientation. Whilst Subjects 1 and 2 are characterised by more focal velocity changes in the brain centre and brainstem, Subject 3 was characterised by higher tissue velocities across the entire brain, most notably during the axial acquisition, indicative of rigid body motion. We anticipate that this led to a considerable perturbation of the DW-SSFP signal from steady-state, leading to substantial signal loss (Supporting Information Figure S6) that impact the performance of parameter estimation. Taken together, these findings motivate improved motion profile initialisation and regularisation to translate the findings presented here to a more generalisable motion-correction DW-SSFP scheme, as described below.

**Figure 13:**
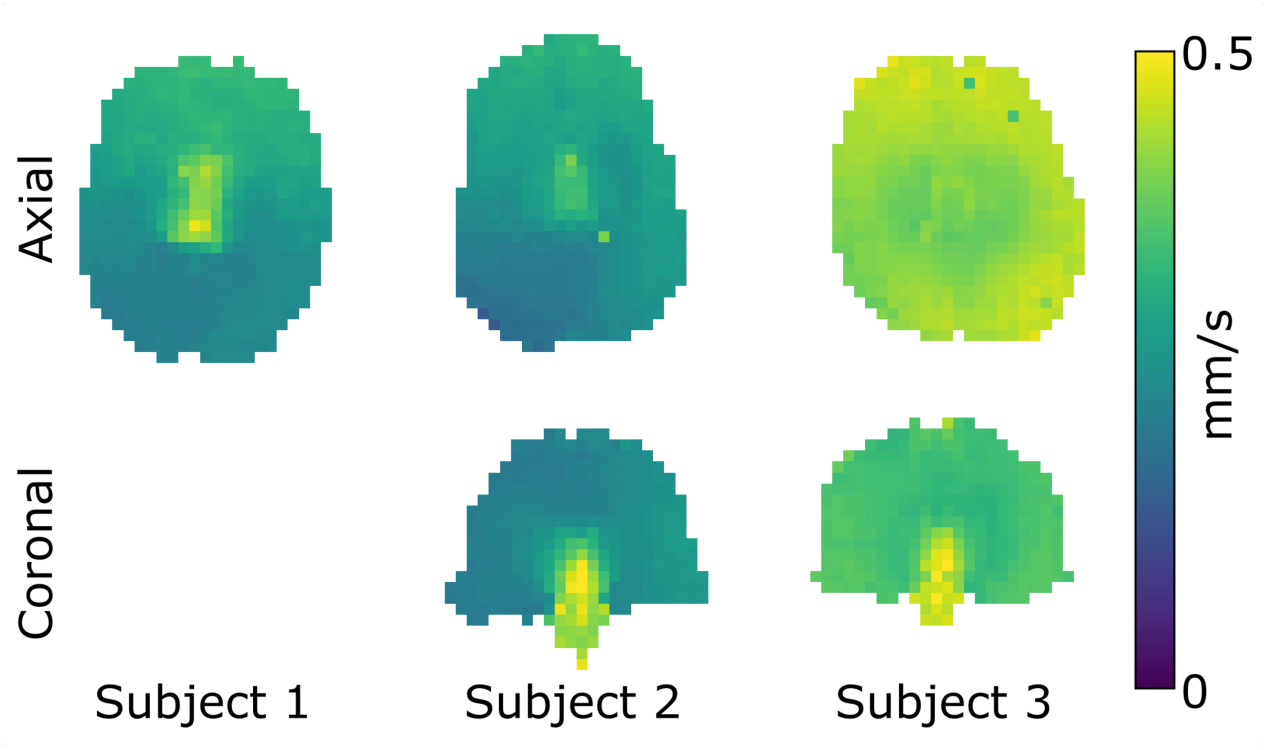
Average absolute velocity along the foot-head gradient orientation. Here we display 𝑉(𝑡) maps representing the average absolute velocity of the timeseries data for all three subjects when the diffusion-gradient was applied along the foot-head orientation. Subject 3 is characterised by a marked increase in the average tissue velocity, indicative of increased subject motion. 𝑉(𝑡) maps estimated from the b100 velocity profiles (Supporting Information Figures S9-S11).

### Translation into a Generalised DW-SSFP Acquisition and Reconstruction Framework

In this work, the proposed EPG-motion framework was applied to 2D, low-resolution single-shot DW-SSFP timeseries data. The acquired data is appropriate for validating the framework and understanding the origin of motion-induced artefacts arising from the signal-forming mechanisms of DW-SSFP, necessitating an EPG (or equivalent) framework to address, with the degree of motion corruption modulated by the local diffusion coefficient (Figure 4 & Supporting Information Figure S4). The simplified fitting approach (non-linear least-squares, single motion estimate per TR, no regularisation) implemented independently for each voxel did not enforce any prior knowledge or constraints on the motion profile, with smooth spatial distributions of 𝑉(𝑡) (Figure 9) arising naturally from the parameter estimation procedure.

When considering a 3D, segmented readout DW-SSFP sequence, the impact of subject motion on reconstructed images will be more severe due to the additional contribution of phase/magnitude inconsistencies between individual k-space segments. To reduce the dimensionality of the reconstruction problem, the proposed framework could be integrated into a motion-reconstruction procedure that performs simultaneous parameter estimation across voxels considering a smoothly varying spatial and temporal motion profile. Pulsatile velocity estimation could initialised with information from complementary methods (e.g. phase contrast imaging^54^, DENSE^52,55^, Amplified MRI^53,56^) and combined with ECG/Pulse oximeter measurements acquired during DW-SSFP acquisition to regularise the fitting procedure. This information could be leveraged with sequences integrating navigators and non-cartesian readouts to establish motion-robust DW-SSFP acquisition schemes.

## Conclusion

We demonstrate that it is possible to model and correct for the impact of subject motion on the magnetisation distribution of DW-SSFP, with motion-corrected experimental DW-SSFP data displaying reduced spatial biases and more consistent contrast with DW-SE EPI tensor estimates. The proposed framework is entirely open source, facilitating future investigations into the impact of motion on DW-SSFP acquisitions.

## Data/Code Availability Statement

The released software (available at github.com/BenjaminTendler/MotionCorrectionDWSSFP, SHA #d0ca7b637a87f96b8c60e3356ce7f9ad85eefc53) provides a framework to investigate the impact of motion on the DW-SSFP signal, allowing for the forward simulation of temporal motion profiles for 1D, 2D and 3D imaging volumes, the resulting DW-SSFP signal, and estimation of motion parameters from DW-SSFP timeseries data. Scripts to replicate many of the figures presented in this manuscript are also provided.

## Acknowledgements

BCT is funded by a Sir Henry Wellcome Postdoctoral Fellowship (Wellcome Trust) [222829/Z/21/Z]. WW is funded by the Royal Academy of Engineering [RF\201819\18\92]. KLM is funded by a Wellcome Trust Senior Research Fellowship [224573/Z/21/Z]. This research was supported by the NIHR Oxford Health Biomedical Research Centre [NIHR203316]. The views expressed are those of the author(s) and not necessarily those of the NIHR or the Department of Health and Social Care. The Centre for Integrative Neuroimaging was supported by core funding from the Wellcome Trust [203139/Z/16/Z and 203139/A/16/Z].

This research was funded in whole, or in part, by the Wellcome Trust [222829/Z/21/Z, 224573/Z/21/Z, 203139/Z/16/Z, 203139/A/16/Z]. For the purpose of open access, the author has applied a CC BY public copyright licence to any Author Accepted Manuscript version arising from this submission.

## Contributions Statement

**BCT** designed the project; established the theory and developed the EPG-motion framework; optimised the DW-SSFP sequence parameters; implemented the acquisition protocol; performed pilot and subject scanning; performed all analysis; wrote the manuscript and produced all figures. **WW** developed the DW-SSFP sequence; and provided manuscript feedback. **KLM** contributed to the project design, providing guidance and feedback on the investigative pathway, theory, and findings; developed the custom DW-SSFP Monte Carlo simulation software; and provided manuscript feedback. **ATH** oversaw the experimental data acquisition, informing experimental design, providing guidance and feedback on the investigative pathway based on experimental pilot data; modified the DW-SSFP sequence to acquire the described experimental data; performed pilot and subject scanning; and provided manuscript feedback.

