## Supporting Information for "Modelling motion-induced signal corruption in steady-state diffusion MRI"

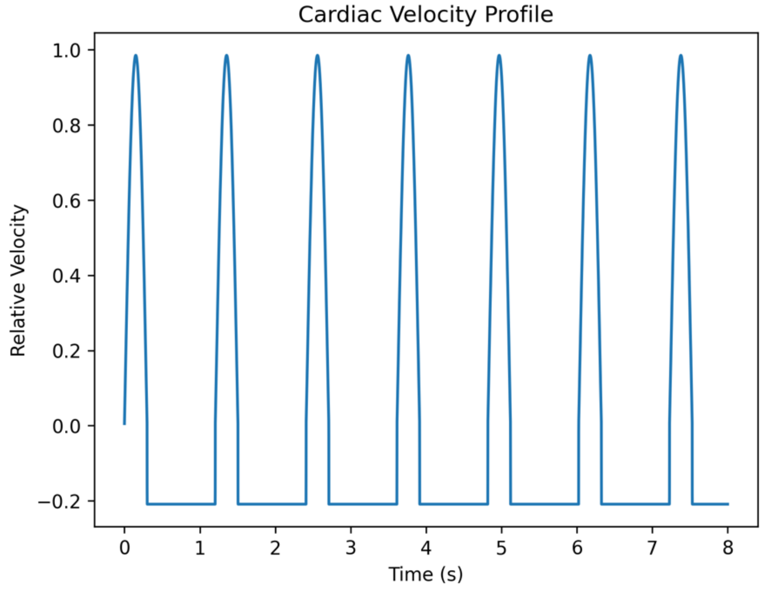


Figure S1: **Pulsatility time-series velocity profile**. Relative velocity timeseries profile was based visually on the experimental thalamus (caudal) profile presented in Greitz et al.[1]. Profile defined modelling systole as a function of $\mathrm{A} \cdot \sin \frac{4\pi \cdot \mathrm{HR} \cdot t}{60}$ over 1/4 of the cardiac cycle, where A = maximum pulsatile velocity (mm/s), HR = heart rate (beats/minute) and $t$ = time. Displacement (i.e. integral of the velocity profile) across a single cardiac cycle was set equal to zero, with the velocity profile defined as constant during diastole. Above profile synthesised setting HR = 50 beats/minute.

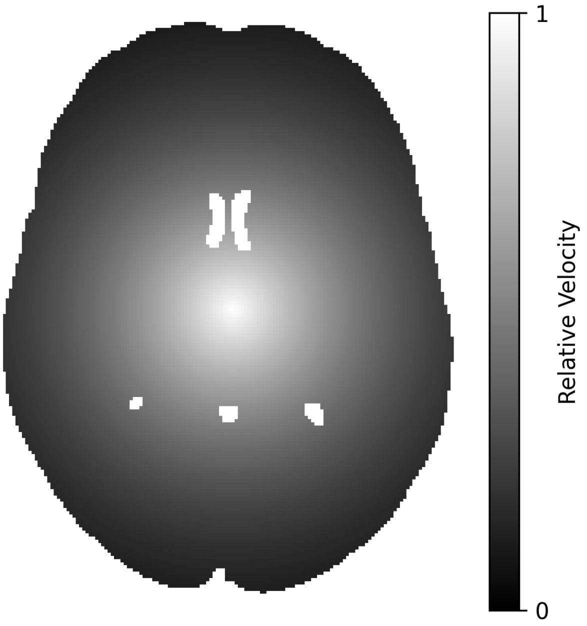


Figure S2: **Pulsatility spatial velocity map**. Relative spatial velocity profile was based visually on the experimental caudal velocity maps presented in Greitz et al.[1] Here, the velocity map displays the relative maximum spatial velocity associated with brain pulsatility, and is used to rescale the cardiac pulsatility time-series profile (Figure S1) in each voxel. Profile defined as a function of $\frac{(-r+\mathrm{c})^3}{\mathrm{c}^3}$, where $r$ = relative voxel position (mm), and c = constant (mm). Here c = 175 mm, preserving the spatial velocity profile for datasets with different voxel sizes or fields of view.

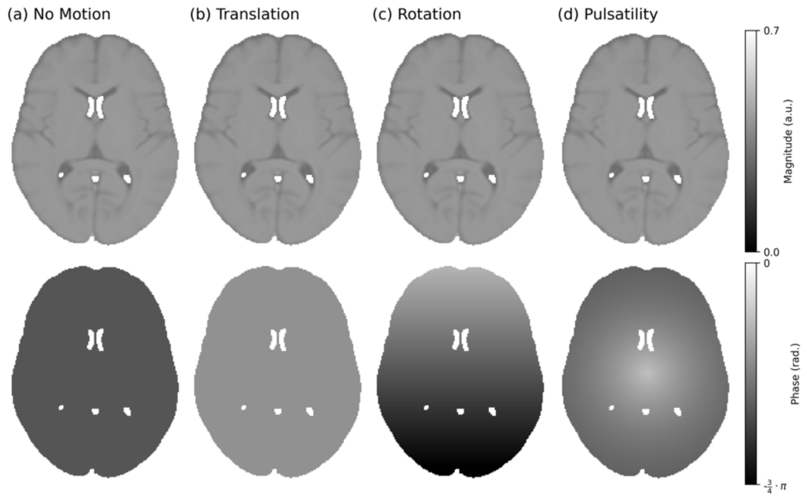


Figure S3: **Impact of motion on DW-SE images**. Here we display simulated single-shot DW-SE images with an instantaneous readout, incorporating (a) no motion (b) constant translation along the foot-head direction ($\vec{v} = [0,0,0.2]$ mm/s), (c) constant rotation with the axis of rotation oriented along the left-right direction (i.e. passing through the sagittal plane) ($\vec{\omega} = [0.2,0,0]$ $^{o}$/s), and (d) cardiac pulsatility along the foot-head direction ($\vec{v}_{\mathrm{Card}} = [0, 0, 0.4]$ mm/s), simulated with the diffusion gradient oriented along the foot-head direction ($\hat{g} = [0,0,1]$). Images synthesised using the HCP1065 DTI template Mean Diffusivity map, available as part of FSL[2]. Cardiac pulsatility timeseries profile and relative velocity map provided in Supporting Information Figures S1 and S2. Sequence parameters based on the experimental investigation performed in Miller & Pauly[3] (defining the TR in DW-SSFP as the DW-SE diffusion time), setting $\mathrm{G} = 40$ mT/m, $\delta = 6.5$ ms, $\alpha = 30^{o}$, $\Delta = 40$ ms, $\phi = 0^{o}$. Relaxation parameters (set equal across all voxels) were based on approximate in vivo values in brain tissue at 3T[4], setting $\mathrm{T}_1 = 832$ ms and $\mathrm{T}_2 = 110$ ms. Voxels were modelled as independent (i.e. the voxel location is stationary over the time-course of the simulation).

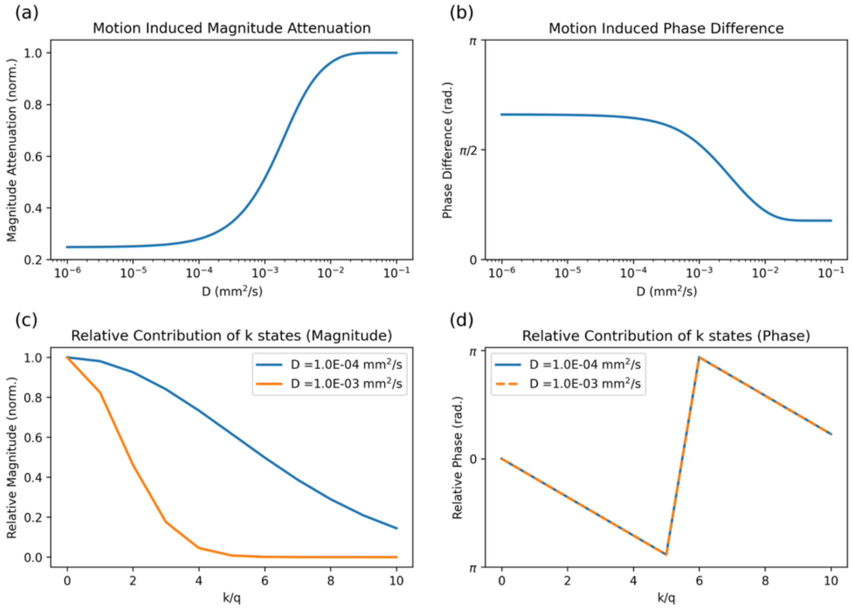


Figure S4: **Interplay between the diffusion coefficient and motion-induced signal changes.** Relative to motion-free simulations, (a) and (b) display the signal amplitude and phase change arising from subject motion as a function of $D$. Higher $D$ leads to a reduced change in signal amplitude and phase, analogous to reduced motion sensitivity. To understand this further, (c) displays the relative diffusion attenuation (log scale) as a function of pathway dephasing order ($k$). Here, increased $D$ (orange line) is associated with a faster loss of the relative signal as a function of $k$. The relative phase difference arising from motion scales linearly as a function of $k_j$, and is independent of $D$ (d). Taken together, tissues with increased $D$ have reduced motion sensitivity arising from the reduced contribution of pathways with high $k$. Motion simulated as a constant translation along a single dimension ($\vec{v} = [0,0,0.2]$ mm/s). Sequence parameters based on the experimental investigation performed in Miller & Pauly[3], setting $\mathrm{G} = 40$ mT/m, $\delta = 6.5$ ms, $\alpha = 30^{o}$, $\mathrm{TR} = 40$ ms, $\phi = 0^{o}$ and $\hat{g} = [0,0,1]$). Relaxation parameters (set equal across all voxels) were based on approximate in vivo values in brain tissue at 3T[4], setting $\mathrm{T}_1 = 832$ ms and $\mathrm{T}_2 = 110$ ms.

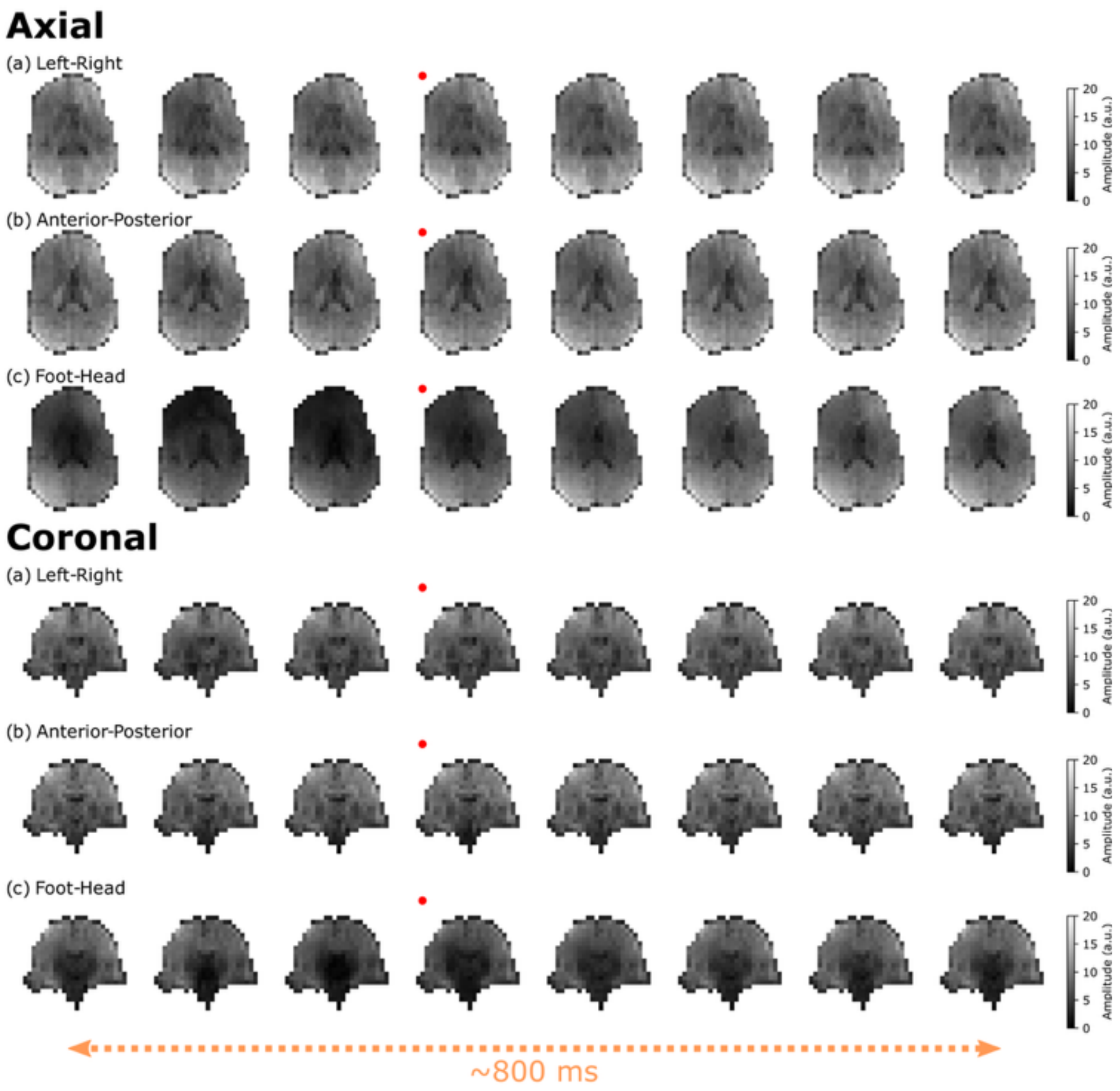


Figure S5: **Experimental DW-SSFP timeseries images (Subject 2).** Equivalent to Figure 8 in the Main Text, (a-c) correspond to example $b_{500}$ timeseries data acquired with left-right (a), anterior posterior (b) and foot-head (c) gradient orientations for an axial (top) and coronal (bottom) slice in Subject 2. The change in signal is most prominent along the foot-head orientation, characterised here by a substantial loss of signal in several brain regions. Red dots indicate the image acquired at the timepoint of the pulse oximeter trigger, with the x-axis reflecting a subset of data acquired over 832 ms, with a gap of 104 ms between images (four TRs). Anterior-posterior images shifted temporally by 54 ms (two TRs) relative to left-right and foot-head orientations. Images have an identical dynamic range to Figure 8 (Main Text).

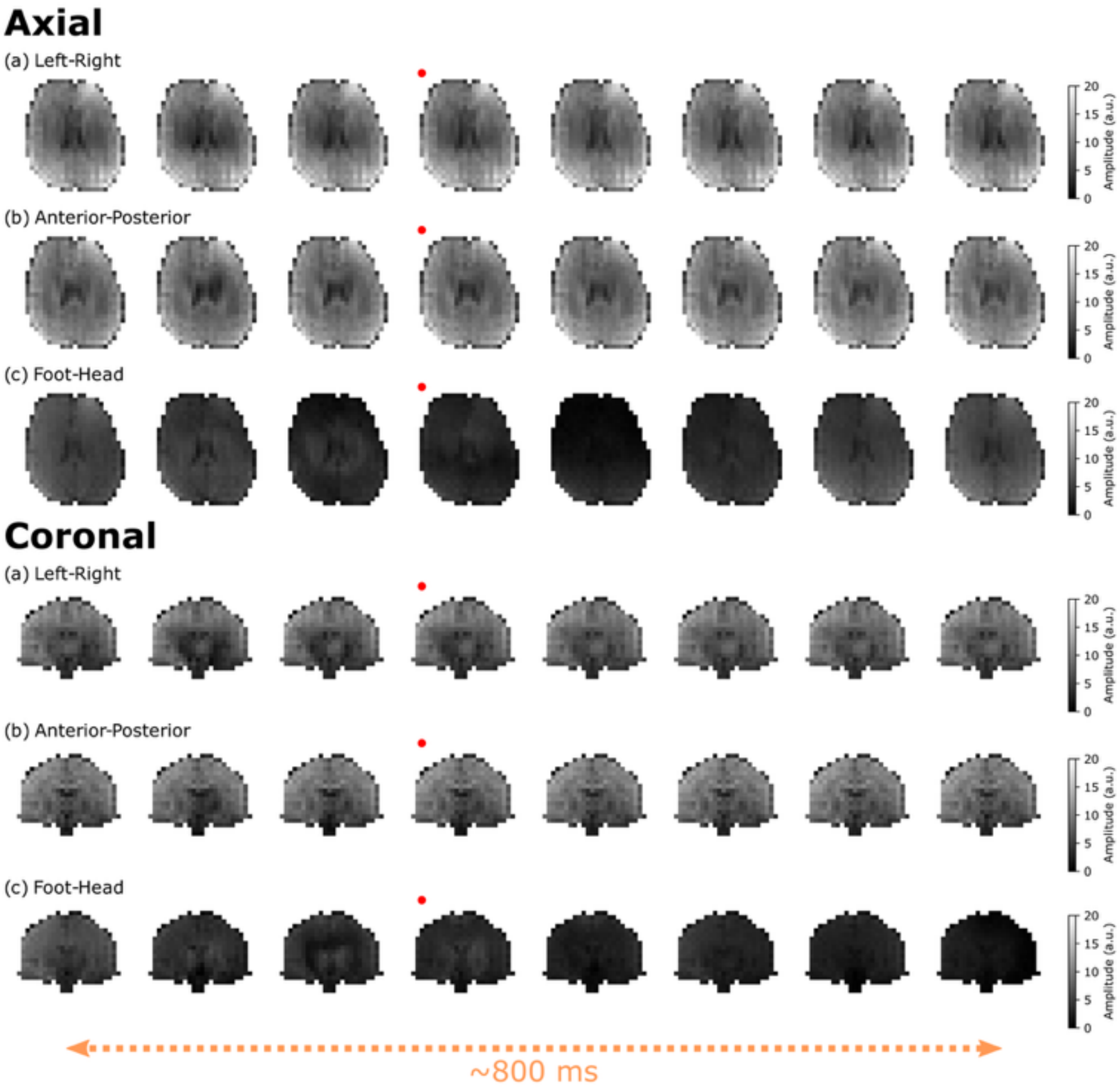


Figure S6: **Experimental DW-SSFP timeseries images (Subject 3).** Equivalent to Figure 8 in the Main Text, (a-c) correspond to example $b_{500}$ timeseries data acquired with left-right (a), anterior posterior (b) and foot-head (c) gradient orientations for an axial (top) and coronal (bottom) slice in Subject 3. The change in signal is most prominent along the foot-head orientation, with an increased level of signal loss in comparison to Subject 1 (Figure 8) and Subject 2 (Supporting Information Figure S5), indicative of increased subject motion. Red dots indicate the image acquired at the timepoint of the pulse oximeter trigger, with the x-axis reflecting a subset of data acquired over 832 ms, with a gap of 104 ms between images (four TRs). Images have an identical dynamic range to Figure 8 (Main Text) and Supporting Information Figure S5.

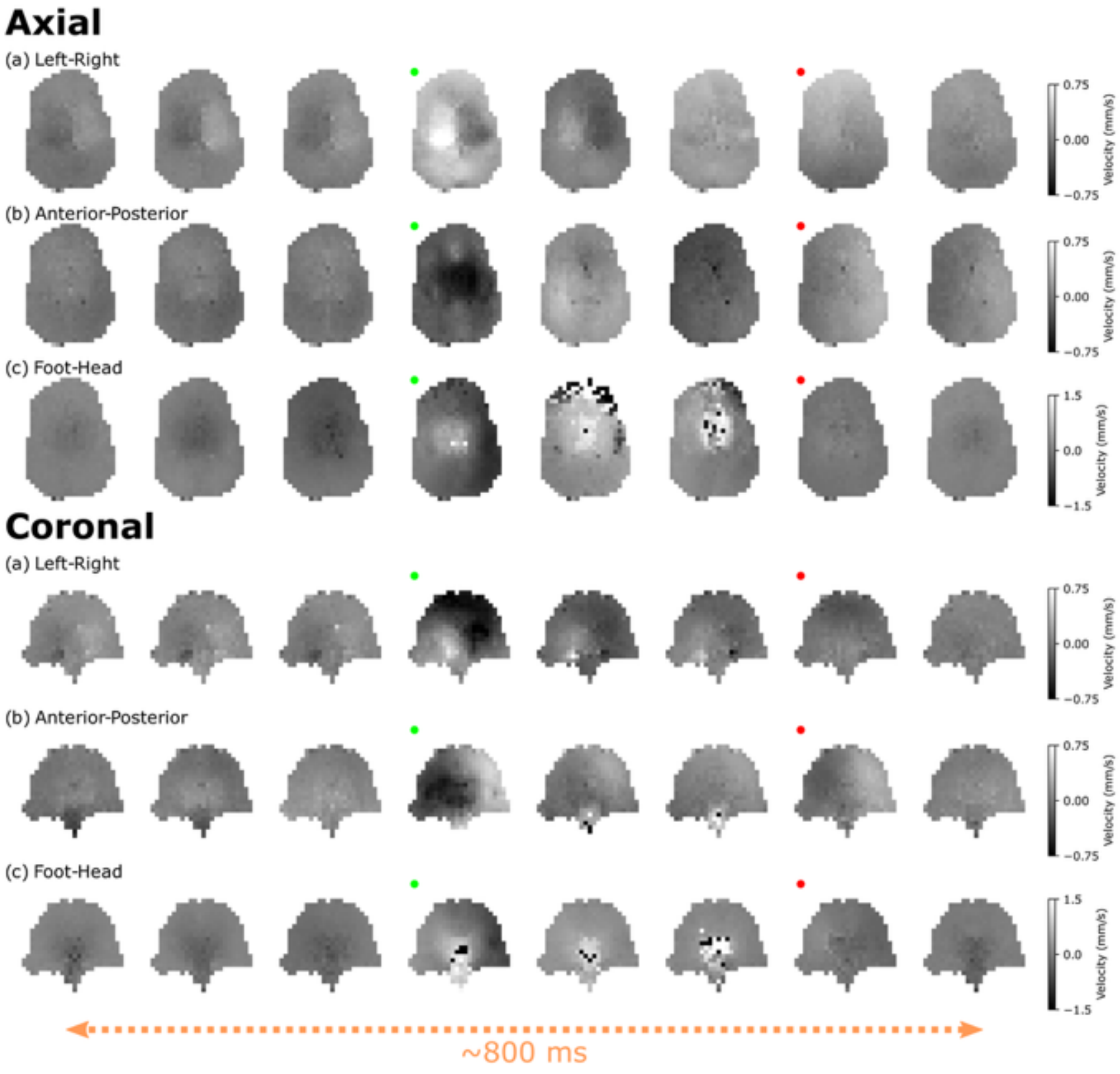


Figure S7: **Estimated $V(t)$ maps (Subject 2).** Equivalent to Figure 9 in the Main Text, (a-c) correspond to example $V(t)$ spatial maps estimated from the experimental $b_{500}$ DW-SSFP data along left-right (a), anterior-posterior (b) and foot-head (c) gradient orientations for an axial (top) and coronal (bottom) slice in Subject 2. Here the red dots indicate the image acquired at the timepoint of the pulse oximeter trigger, with the green dots indicating peak brain tissue velocity changes. The x-axis reflects a subset of data acquired over 832 ms, with a gap of 104 ms between images (four TRs). Anterior-posterior images shifted temporally by 54 ms (two TRs) relative to left-right and foot-head orientations. $V(\mathrm{t})$ defined as the component of instantaneous velocity in the direction of the diffusion gradient across TRs (Eq. [4]). Black/white voxels within the brain correspond to velocities exceeding the colour bar limits (right hand side). Velocity maps have an increased dynamic range in comparison to Figure 9 (Main Text).

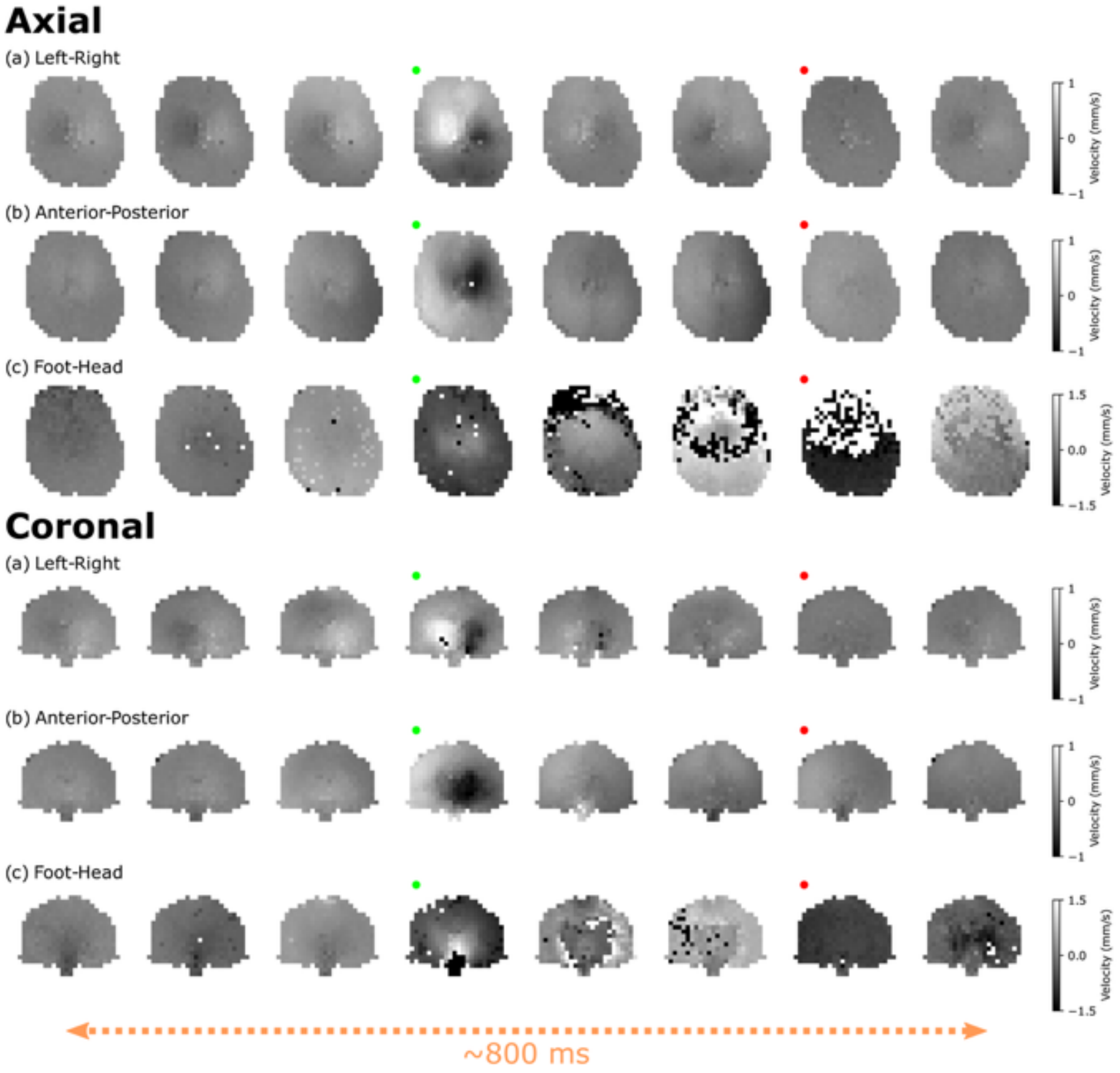


Figure S8: **Estimated $V(t)$ maps (Subject 3).** Equivalent to Figure 9 in the Main Text, (a-c) correspond to example $V(t)$ spatial maps estimated from the experimental $b_{500}$ DW-SSFP data along left-right (a), anterior-posterior (b) and foot-head (c) gradient orientations for an axial (top) and coronal (bottom) slice in Subject 3. Here the red dots indicate the image acquired at the timepoint of the pulse oximeter trigger, with the green dots indicating peak brain tissue velocity changes. The x-axis reflects a subset of data acquired over 832 ms, with a gap of 104 ms between images (four TRs). $V(\mathrm{t})$ defined as the component of instantaneous velocity in the direction of the diffusion gradient across TRs (Eq. [4]). Black/white voxels within the brain correspond to velocities exceeding the colour bar limits (right hand side). Velocity maps have an increased dynamic range in comparison to Figure 9 (Main Text) (all gradient orientations) and Supporting Information Figure S7 (left-right and anterior-posterior gradient orientations).

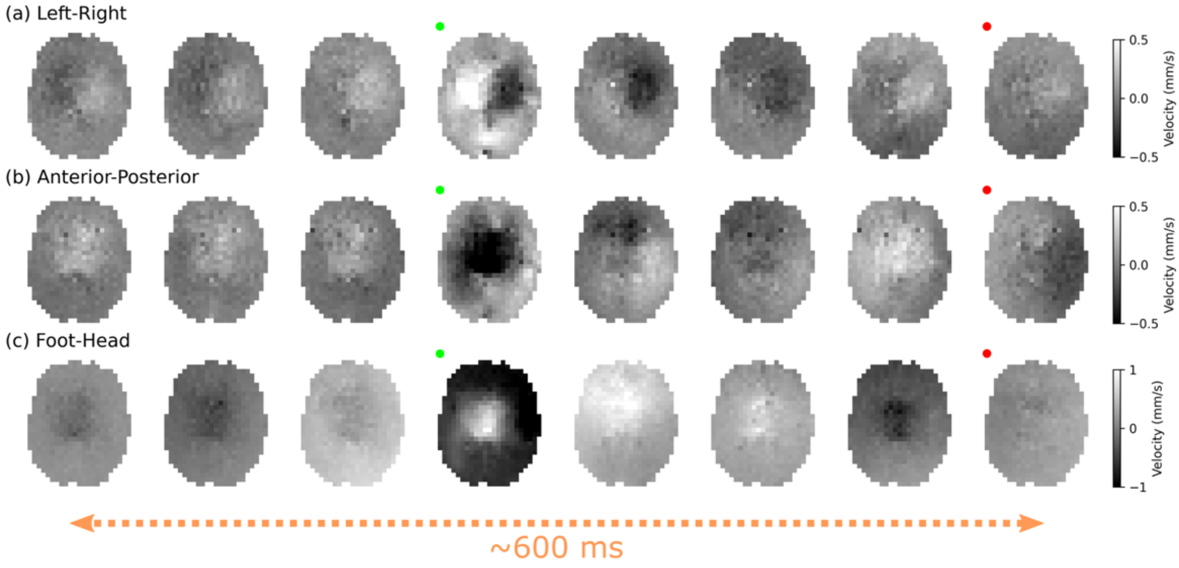


Figure S9: **Estimated $b_{100}$ $V(t)$ maps (Subject 1).** (a-c) correspond to $V(t)$ maps estimated from the experimental DW-SSFP $b_{\text{eff}} = 100 \text{ s/mm}^2$ data along the left-right (a), anterior posterior (b), and foot-head (c) gradient orientations over the cardiac cycle. The displayed maps are equivalent to the $b_{500}$ maps displayed in Figure 9 (Main Text), where $V(t)$ maps are more consistently estimated along the foot-head direction. Here the red dots indicate the image acquired at the timepoint of the pulse oximeter trigger, with the green dots indicating peak brain tissue velocity changes. The x-axis reflects a subset of data acquired over 624 ms, with a gap of 78 ms between images (three TRs). $V(\text{t})$ defined as the component of instantaneous velocity in the direction of the diffusion gradient across TRs (Eq. [4]). Black/white voxels within the brain correspond to velocities less/greater than the colour bar limits (right hand side). Corresponding figures for Subjects 2 and 3 are provided in Supporting Information Figures S10 and S11.

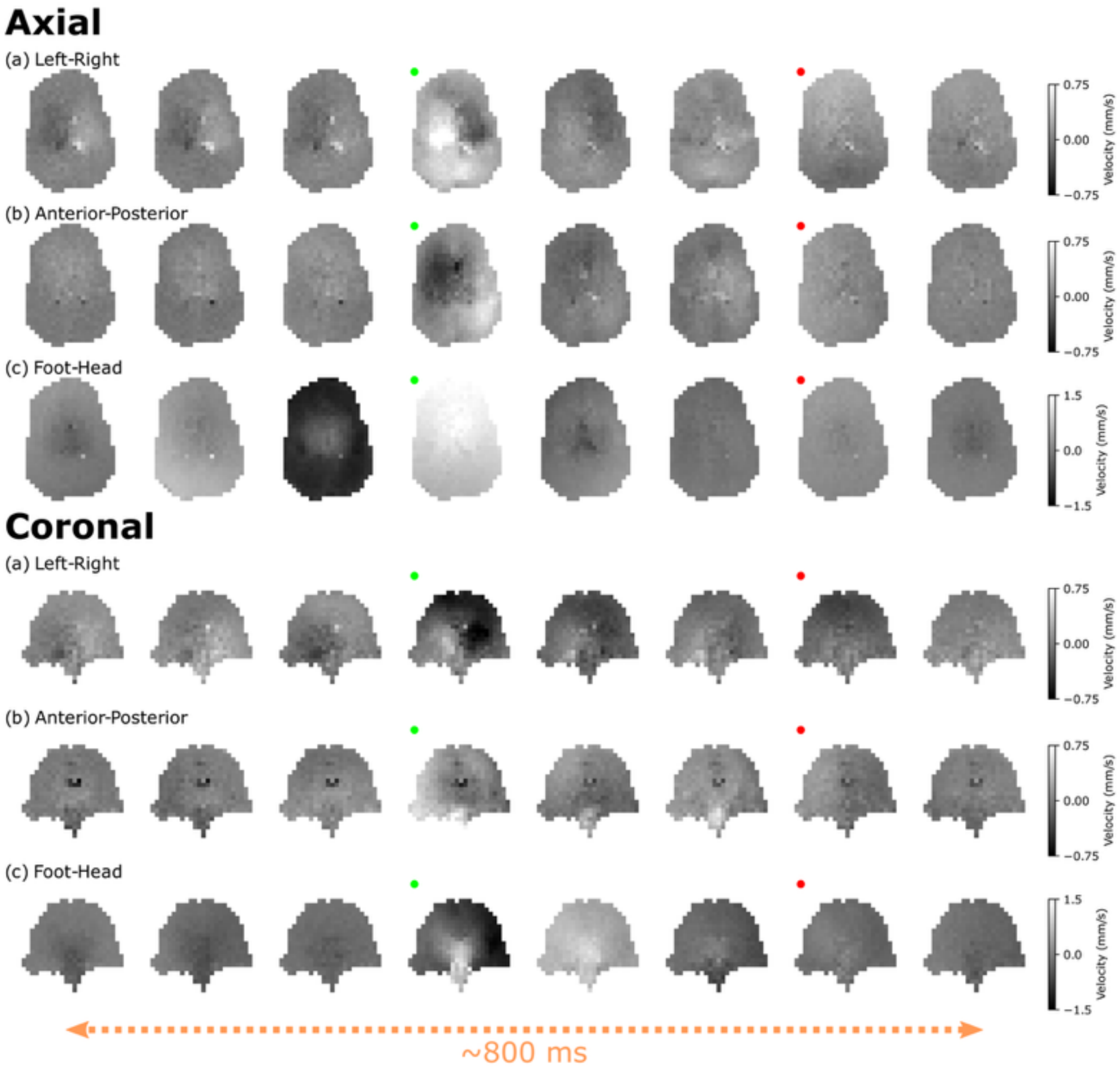


Figure S10: **Estimated $b_{100}$ $V(t)$ maps (Subject 2).** Equivalent to Supporting Information Figure S9, here we display $V(t)$ maps estimated from the experimental DW-SSFP $b_{\mathrm{eff}} = 100\ \mathrm{s/mm^2}$ data along the left-right (a), anterior posterior (b), and foot-head (c) gradient orientations over the cardiac cycle for an axial (top) and coronal (bottom) slice in Subject 2. The displayed maps are equivalent to the $b_{500}$ maps displayed in Supporting Information Figure S7, where $V(t)$ maps are more consistently estimated along the foot-head direction. Here the red dots indicate the image acquired at the timepoint of the pulse oximeter trigger, with the green dots indicating peak brain tissue velocity changes. The x-axis reflects a subset of data acquired over 832 ms, with a gap of 104 ms between images (four TRs). $V(\mathrm{t})$ defined as the component of instantaneous velocity in the direction of the diffusion gradient across TRs (Eq. [4]). Black/white voxels within the brain correspond to velocities less/greater than the colour bar limits (right hand side). Velocity maps have an increased dynamic range in comparison to Supporting Information Figure S9.

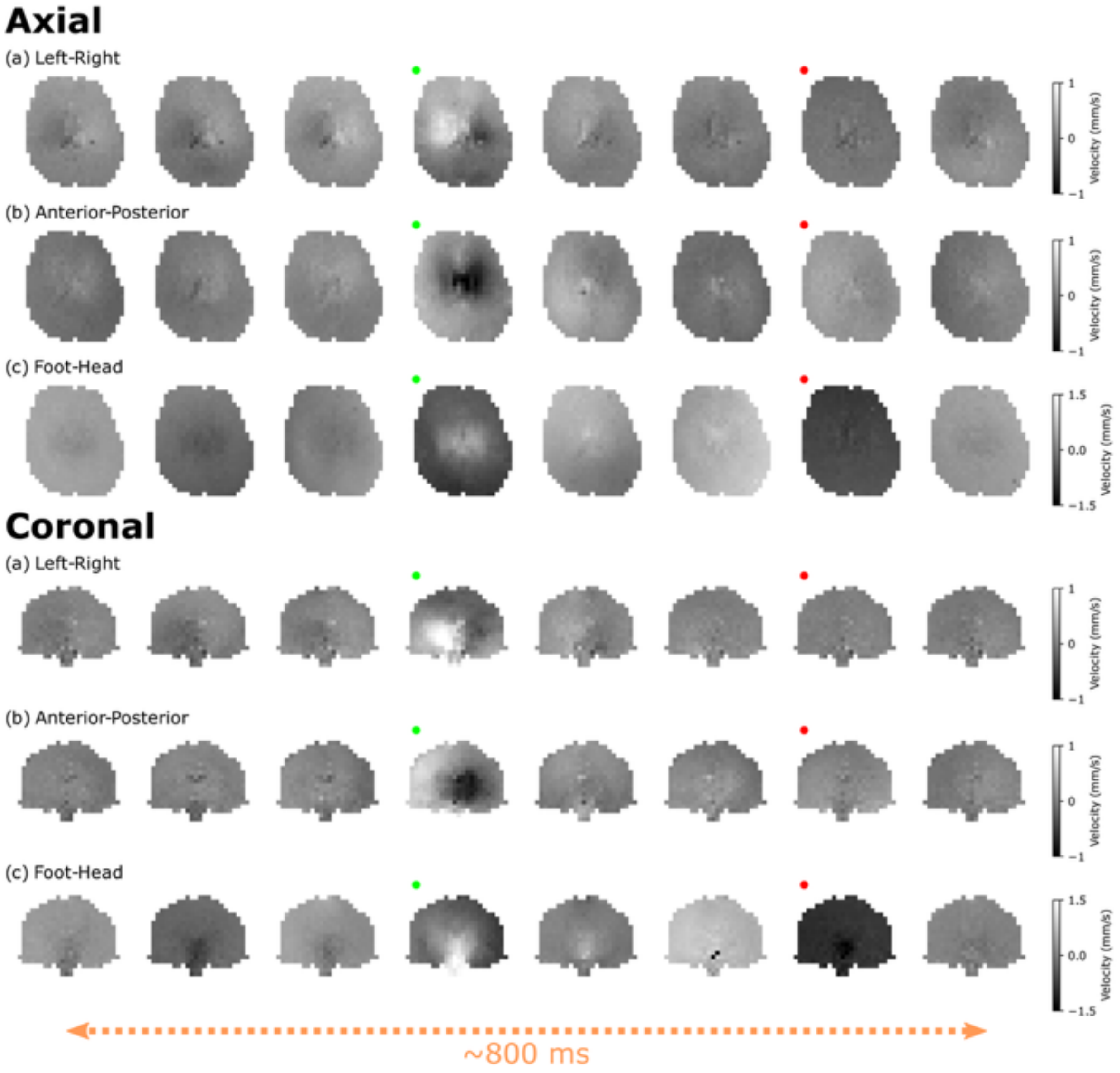


Figure S11: **Estimated $b_{100}$ $V(t)$ maps (Subject 3).** Equivalent to Supporting Information Figures S9 and S10, here we display $V(t)$ maps estimated from the experimental DW-SSFP $b_{\text{eff}} = 100\ \text{s/mm}^2$ data along the left-right (a), anterior posterior (b), and foot-head (c) gradient orientations over the cardiac cycle for an axial (top) and coronal (bottom) slice in Subject 2. The displayed maps are equivalent to the $b_{500}$ maps displayed in Supporting Information Figure S8, where $V(t)$ maps are more consistently estimated along the foot-head direction. Here the red dots indicate the image acquired at the timepoint of the pulse oximeter trigger, with the green dots indicating peak brain tissue velocity changes. The x-axis reflects a subset of data acquired over 832 ms, with a gap of 104 ms between images (four TRs). $V(\text{t})$ defined as the component of instantaneous velocity in the direction of the diffusion gradient across TRs (Eq. [4]). Black/white voxels within the brain correspond to velocities less/greater than the colour bar limits (right hand side). Velocity maps have an increased dynamic range in comparison to Supporting Information Figure S9 (all gradient orientations) and Supporting Information Figure S10 (left-right and anterior-posterior gradient orientations).

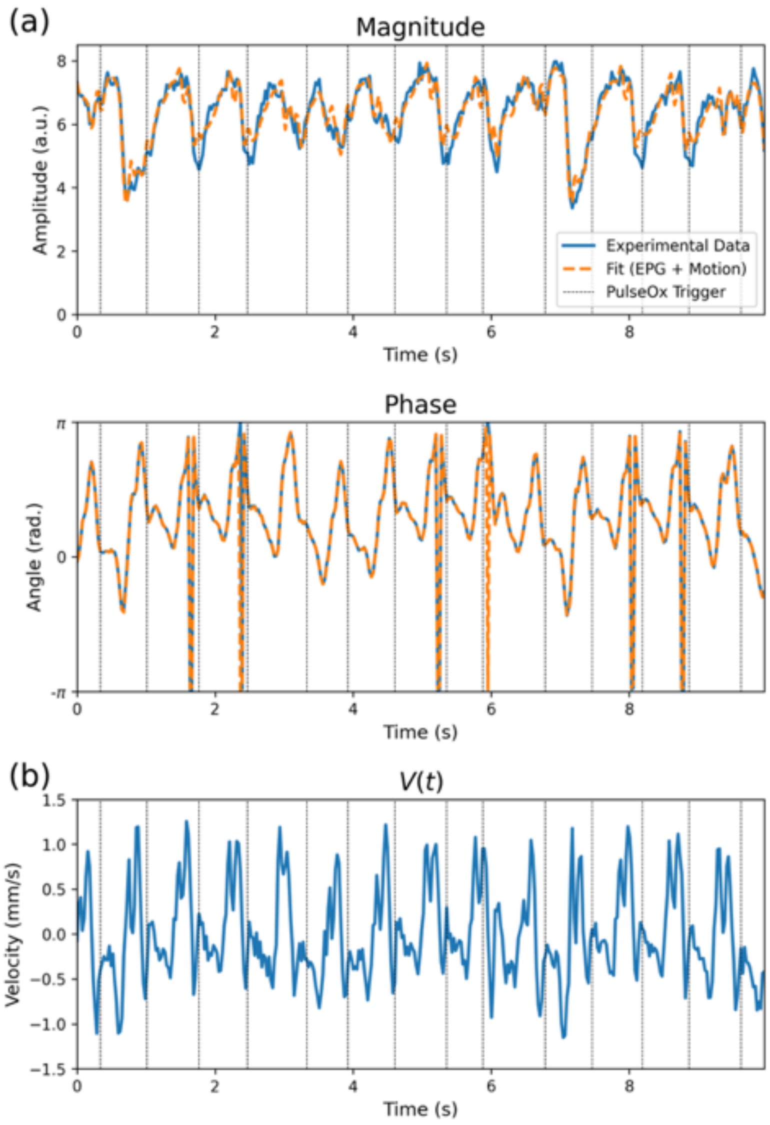


Figure S12: **DW-SSFP signal and $V(t)$ profile in a single voxel of experimental $b_{100}$ data**. Identical in structure to Figure 7 (Main Text), here we display an experimental (blue lines) and estimated (orange lines) DW-SSFP signal profile (a) and the $V(t)$ profile (b) for a single voxel of $b_{eff} = 100 \text{ s/mm}^2$ data acquired in the thalamus (visualised in inset figure of experimental data in (a) – red voxel), with the diffusion gradient oriented along the foot-head axis. Black vertical lines correspond to pulse oximeter trigger timings acquired as part of the acquisition. $V(t)$ does not exceed 1.5 mm/s (maximum 1.26 mm/s), with the change in velocity per TR below 1 mm/s (maximum 0.86 mm/s/TR). DW-SSFP signal multiplied by $e^{-i}$ to facilitate visualisation of wrapped phase data.

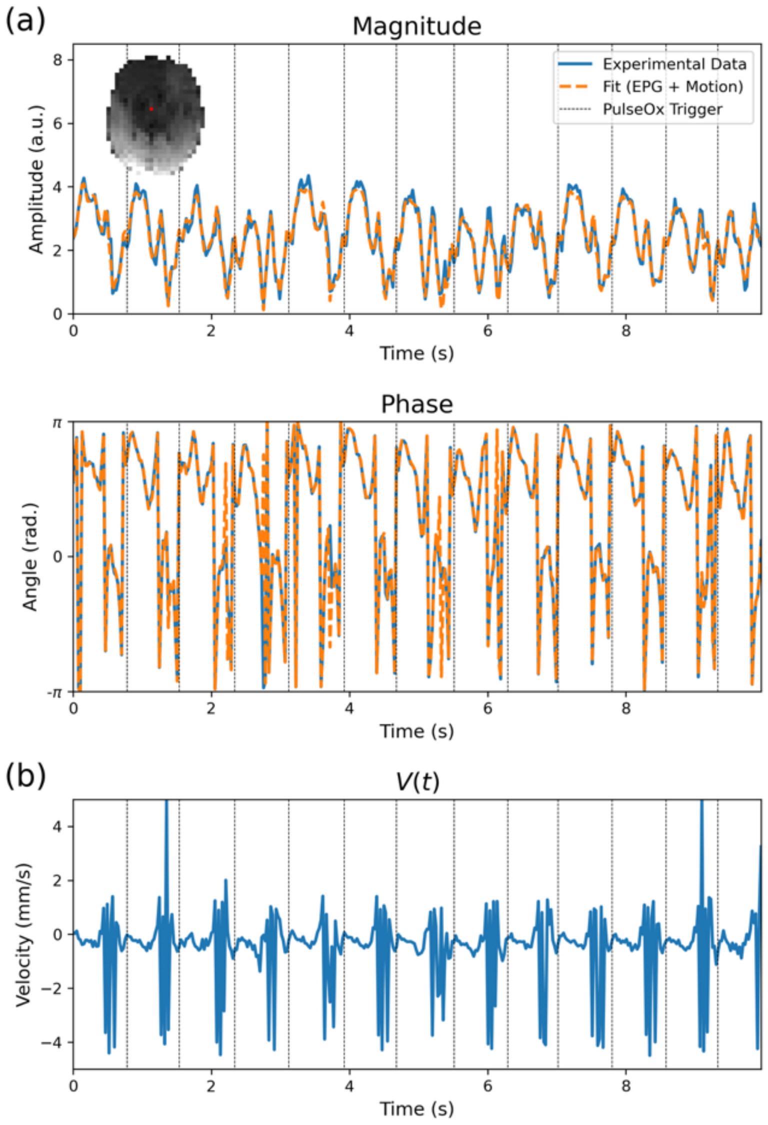


Figure S13: **DW-SSFP signal and $V(t)$ profile in a single voxel of experimental $\mathbf{b_{eff}} = \mathbf{500\ s/mm^2}$ data**. Identical in structure to Figure S12, here we display an experimental (blue lines) and estimated (orange lines) DW-SSFP signal profile (a) and the $V(t)$ profile (b) for a single voxel of $b_{500}$ data acquired in the thalamus (visualised in inset figure of experimental data in (a) – red voxel), with the diffusion gradient oriented along the foot-head axis. Black vertical lines correspond to pulse oximeter trigger timings acquired as part of the acquisition. $V(t)$ estimates a consistent rapidly oscillating velocity profile during systole with velocity values routinely exceeding 3 mm/s, far greater than the equivalent velocity profile estimated from the $b_{100}$ data (Figure S12). DW-SSFP signal multiplied by $e^{-i}$ to facilitate visualisation of wrapped phase data.

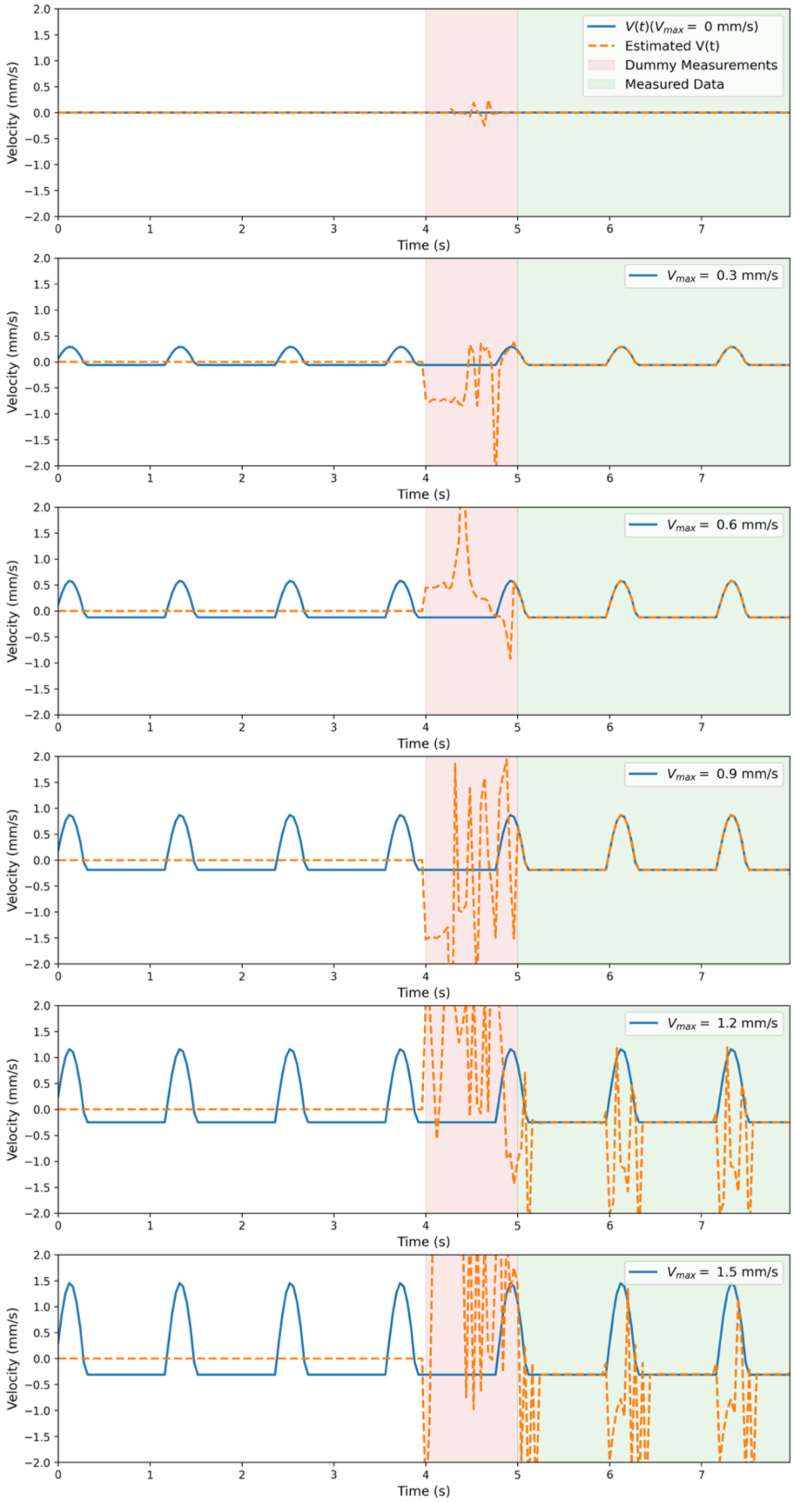


Figure S14: **Simulated and estimated $V(t)$ profiles.** Here we display the simulated (blue) and estimated (orange) $V(t)$ profiles ($V_{max}$ ranging from 0 to 1.5 mm/s) from noiseless data, with sequence parameters based on the experimental investigation performed in Miller & Pauly[3], setting $G = 40$ mT/m, $\delta = 6.5$ ms, $\alpha = 30^o$, TR $= 40$ ms and $\phi = 0^o$. Excellent agreement is given over the $V(t)$ profiles for velocities below 1 mm/s, with errors in velocity estimation during systole at 1.2 and 1.5 mm/s. Despite the discrepancy during systole, the diffusion coefficient can still be reliably estimated (Figure 6). Velocity estimation during systole could be improved with more robust initialisation of the velocity profile based on expected pulsatility waveforms or complementary data (see Discussion). $V(\mathrm{t})$ defined as the component of instantaneous velocity in the direction of the diffusion gradient across TRs (Eq. [4]).

| | $D$ (mm$^2$/s) | $\phi$ (rad) | $V(t)$ (mm/s) |
|---|---|---|---|
| Initialisation | $0.5 \cdot 10^{-3}$ | $\pi/2$ | $[0,0,0,\ldots]$ |
| Lower Bound | $0$ | $-20\pi$ | $[-1.5,-1.5,-1.5,\ldots]$ |
| Upper Bound | $10 \cdot 10^{-3}$ | $20\pi$ | $[1.5,1.5,1.5,\ldots]$ |

Supporting Information Table S1: **Initialisation parameters and bounds for simulated data fitting.**

| **ep2d_se ($T_1$)** | | **se_mc ($T_2$)** | |
|---|---|---|---|
| Resolution | 3.4 mm iso. | Resolution | 3.4 mm iso. |
| Matrix size | 64x64x1 | Matrix size | 64x64x1 |
| No Inversions | 12 | No Echoes | 32 |
| TE | 48 ms | TEs | 14.3:14:3:457.6 |
| TR | 15 s | TR | 5 s |
| Flip Angle | 90°/180° | Flip Angle | 90°/180° |
| Bandwidth | 1660 Hz/Pixel | Bandwidth | 115 Hz/Pixel |
| TIs | 100, 200, 310, 440, 580, 750, 940, 1170, 1470, 1890, 2580, 4980 ms | | |
| **3DREAM**[5] | | **Diffusion-Weighted Spin Echo Echo Planar Imaging (DW-SE EPI)** | |
| Resolution | 6.5 mm iso. | Resolution | 3x3x6 mm$^3$ |
| Matrix size | 34x34x40 | Matrix size | 64x64x1 |
| TEs | 11, 21 ms | b-value | 0, 500 s/mm$^2$ |
| TR | 6 s | no. Directions | 12 |
| Flip angle | 60° | no. b0s | 1 |
| Bandwidth | 1000 Hz / Pixel | TE | 69 ms |
| | | TR | 5 s |
| | | Flip Angle | 90°/180° |
| | | Bandwidth | 2365 Hz/Pixel |

Supporting Information Table S2: **Sequence parameters for supporting acquisitions.** $T_1$ map estimated from the ep2d_se data assuming monoexponential signal evolution (custom code). $T_2$ map estimated from the se_mc data using via an EPG model (custom code), as described in Tendler et al.[6] $B_1$ map estimated from 3DREAM[5] sequence provided by automated scanner reconstruction. The DW-SE EPI data were first downsampled to the equivalent resolution of the DW-SSFP data (32x32), with diffusion coefficients subsequently estimated for each diffusion gradient orientation and reconstructed into a tensor using custom code. Datasets were coregistered to the DW-SSFP data prior to processing using FLIRT[7,8].

| | $D$ (mm$^2$/s) | $\phi_{100}/\phi_{500}$ (rad) | $V_{100}/V_{500}(t)$ (mm/s) |
|---|---|---|---|
| Initialisation | $0.5 \cdot 10^{-3}$ | $\pi/2$ | $[0,0,0,\ldots]$ |
| Lower Bound | $1 \cdot 10^{-6}$ | $-20\pi$ | $[-5,-5,-5,\ldots]$ |
| Upper Bound | $10 \cdot 10^{-3}$ | $20\pi$ | $[5,5,5,\ldots]$ |

Supporting Information Table S3: **Initialisation parameters and bounds for experimental data fitting.**
